# Secondary metabolites of *Bacillus subtilis* impact soil-derived semi-synthetic bacterial community assembly

**DOI:** 10.1101/2020.08.20.259788

**Authors:** Heiko T. Kiesewalter, Carlos N. Lozano-Andrade, Mikael L. Strube, Ákos T. Kovács

## Abstract

Secondary metabolites provide *Bacillus subtilis* with increased competitiveness towards other microorganisms. In particular nonribosomal peptides (NRPs) have an enormous antimicrobial potential by causing cell lysis, perforation of fungal membranes, enzyme inhibition or disruption of bacterial protein synthesis. This knowledge was primarily acquired *in vitro* when *B. subtilis* was competing with other microbial monocultures. However, our understanding of the true ecological role of these small molecules is limited.

In this study, we have established soil-derived semi-synthetic mock communities containing 13 main genera and supplemented them with *B. subtilis* P5_B1 WT, its NRP deficient strain *sfp* or single NRP mutants incapable of producing surfactin, plipastatin or bacillaene. 16S amplicon sequencing revealed that the invasion of NRP-producing *B. subtilis* strains had no major impact on the bacterial communities. Still, the abundances of the two genera *Lysinibacillus* and *Viridibacillus* were reduced. Interestingly, this effect was diminished in communities supplemented with the NRPs deficient strain. Growth profiling of *Lysinibacillus fusiformis* M5 exposed to either spent media of the *B. subtilis* strains or pure surfactin indicates the sensitivity of this strain towards the biosurfactant surfactin.

Our study provides a more in-depth insight into the influence of *B. subtilis* NRPs on semi-synthetic bacterial communities and helps to understand their ecological role.

## Introduction

In nature, bacteria live in complex communities where they interact with various other microorganisms. Most microbial communities are influencing biochemical cycles and impact agriculture, from which the latter is primarily mediated due to plant growth promotion [1]– [4]. Extensive research has been conducted in the last decade to scrutinise the occurring natural processes and their impact on the environment, investigate the functions and interactions of community members such as metabolite cross-feeding interactions, and to engineer them eventually [5]–[7]. The soil is one of the five main habitats on Earth of bacteria and archaea [8]. It is very heterogeneous since it exhibits spatial variability in nutrient availability and geochemical feature [9]. Therefore, the soil consists of microbial hotspots indicating faster process rates than the average soil [10]. One such microbial hotspot is the rhizosphere harbouring microbial communities where various interactions between bacteria, fungi and plants take place [11]. The composition of microbial communities depends on multiple factors. Studies have revealed that the composition of bacterial soil communities varied during different seasons at the same sampling sites [12],[13]. Moreover, it has been recently demonstrated that precipitation rates had a significant impact on bacterial communities since bacterial soil communities had a higher diversity in dry than in rainy seasons [14]. Besides the seasonal factors, even different plant species with their varying root exudates as well as various soil types impact the microbial community composition in the rhizosphere [15]–[20]. Microbial communities can consist of hundreds and thousands of diverse species, which makes investigations very challenging and hard to reproduce. One alternative approach is to establish a host-associated synthetic community, usually with members from the same kingdom, with a defined composition but fewer members [19],[21]. Lebeis *et al.* used an artificial community of 38 bacterial strains to demonstrate that plant phytohormones sculpt the root microbiome [19]. In comparison, Niu *et al.* established a seven-species bacterial community based on host selection to mimic the principle root microbiome of maise [22].

An important mediator of interactions between microorganisms is believed to be secondary metabolites (SMs) [23]. Many of them are well-studied *in vitro*, but the true ecological role of SMs is still subject of investigation. Different opinions about their primary role in nature exist in the literature, some share the view that SMs are mainly microbial weapons, but others instead designate them as signalling molecules [24]–[27]. Additionally, Pettit (2009) and Wakefield *et al.* (2017) have demonstrated that some bacterial or fungal biosynthetic gene clusters are silent when strains are grown in monocultures under standard laboratory conditions but are expressed in intra- or inter-kingdom co- or multi-cultures [28],[29]. Furthermore, they could show that some SMs had a higher production rate in multi-cultures, highlighting that neighbouring organisms induce and increase SM production in the tested strains.

*Bacillus subtilis* is a well-studied soil bacterium and is used as a model organism for biofilm formation and sporulation [30]. It has been shown that several members of the *B. subtilis* species complex have exceptional plant-growth-promoting and plant-health-improving properties by suppressing plant pathogenic bacteria and fungi [31]. However, it is not completely understood how soil administered *Bacillus* spp. affect the indigenous microbial communities. Gadhave *et al.* have shown that supplementation of *B. subtilis, Bacillus amyloliquefaciens* (now identified as *Bacillus velezensis*) and *Bacillus cereus* to the roots of broccoli plants led to species-dependent changes in the diversity, evenness, and relative abundances of endophytic bacterial communities [32]. Like many other soil bacteria, *B. subtilis* and other *Bacillus* spp. produce various SMs [33],[34]. The most prominent and bioactive SMs are nonribosomal peptides (NRPs), whose isoforms belong to the families of surfactins, fengycins or iturins [35],[36]. They are biosynthesised by large enzyme complexes, nonribosomal peptide synthetases (NRPS). For the biosynthesis of *B. subtilis* NRPs, the phosphopantetheinyl transferase *Sfp* is needed since it has been shown to activate the peptidyl carrier protein domains, converting it from the inactive apo-form to the active holo-form [37]. *B. subtilis* has four *sfp*-dependent SMs, of which three are synthesised by NRPS gene clusters (surfactin, plipastatin and bacillibactin) and one by a hybrid NRPS-PKS gene cluster (bacillaene). The well-studied biosurfactant surfactin, encoded by the *srfAA-AD* gene cluster, reduces the surface tension needed for swarming and sliding motility [38],[39]. Its bioactivity is specifically evoked by the surfactant activity triggering cell lysis due to penetration of bacterial lipid bilayer membranes and formation of ion-conducting channels [40]–[42]. The bioactivity of surfactin was shown against *Listeria* spp. and *Legionella monocytogenes* [43],[44]. It is presumed that the antifungal plipastatin, expressed from the *ppsA-E* gene cluster, acts as an inhibitor of phospholipase A2, forming pores in the fungal membrane and causing morphological changes in fungal membrane and cell wall [45],[46]. Its antifungal potential primarily against various filamentous fungi was demonstrated [47]–[51]. The broad-spectrum antibiotic bacillaene, synthesised by the *pksB-S* gene cluster, is mainly targeting bacterial protein synthesis [52]. Still, it was also shown that it could protect cells and spores from predation [53]. We recently demonstrated that the production of these NRPs varies among co-isolated *B. subtilis* environmental strains due to missing core genes or potentially altered gene regulation highlighting the existing natural diversity of SM production in this species [51].

In this study, we focus on soil-derived semi-synthetic bacterial mock communities and describe how these are affected by a *B. subtilis* strain that was previously isolated from the same sampling site from which the bacterial mock communities originated. We investigated with an NRP mutant-based approach the impact of NRPs on the establishment and composition of the bacterial communities. We previously demonstrated that strain P5_B1 produces the NRPs surfactin and plipastatin and has further BGC predictions for the NRPs bacillaene and bacillibactin [51]. 16S rRNA amplicon sequencing revealed that the established semi-synthetic mock communities contained 13 genera with a relative abundance > 0.19 % in at least one mock community. Furthermore, it demonstrated that the addition of *B. subtilis* suppressed the genera *Lysinibacillus* and *Viridibacillus*. Additional optical density (OD)-based growth monitoring of the selected strain *Lysinibacillus fusiformis* M5 confirmed the impact of *B. subtilis*-produced surfactin on its growth.

## Results

### Impact of *B. subtilis* secondary metabolites on taxonomic groups in semi-synthetic mock communities

We established soil-derived semi-synthetic mock communities and supplemented them with *B. subtilis* WT P5_B1, its NRP deficient strain (*sfp*) or its single NRP mutants incapable of producing either surfactin (*srfAC*), plipastatin (Δ*ppsC*) or bacillaene (Δ*pksL*), or kept untreated culture as control (Figure 1). To investigate the impact of *B. subtilis* NRPs on the bacterial community composition, we sequenced and analysed amplicons of the V3-V4 region of the 16S rRNA gene. The taxonomic summaries give an overview of the relative abundance of the most frequent genera present in each assay and replicate (Figure 2). We investigated the taxonomic level genus since we could not observe any differences among the treated and untreated communities at the class level and similar observations between the family and genus levels. Moreover, the targeted V3-V4 region of the 16S rRNA gene does not allow sufficient distinction below this rank. Unsurprisingly, the two soil samples differ tremendously from the *in vitro* samples and indicating a higher genus richness. We determined that *Bacillus* was the most abundant genus in the two soil samples with a relative proportion between 19 and 35 %. Other genera with abundances higher than 2 % were *Sporosarcina* (4 – 11 %), *Candidatus* Udaeobacter (7 – 10 %) and *Gaiella* (3 – 4 %). The communities of the 12 h-pre-cultivated soil suspension consisted primarily of the two genera *Bacillus* (56 – 65 %) and *Acinetobacter* (29 – 34 %). Additional genera with an abundance higher than 1 % were *Lysinibacillus* (1.2 – 3.2 %), *Pseudomonas* (1.0 – 2.2 %) and *Viridibacillus* (0.6 – 1.5 %). The genus richness of the four pre-cultured soil suspensions was between 12 and 18 of in total 21 genera.

**Figure 1.**
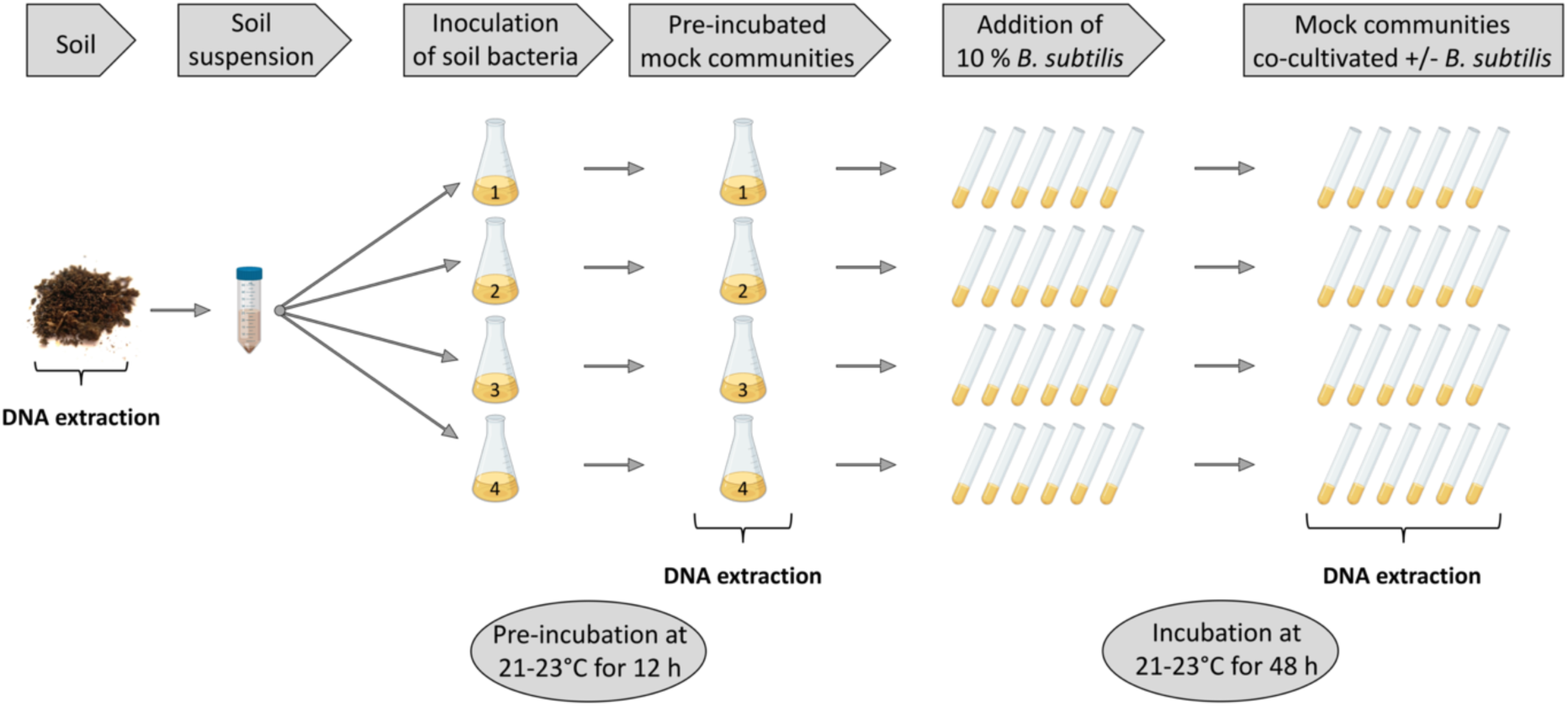
Overview of the experimental setup. A soil suspension, obtained from a soil sample, was used as inoculum for four independent replicates and pre-incubated for 12 h. Enriched pre-cultures were aliquoted and supplemented with 10 % *B. subtilis* strains or left untreated and incubated for 48 h. DNA was extracted from the soil sample, pre-incubated soil suspensions and mock communities. Part of Figure 1 has been created using BioRender.com.

**Figure 2.**
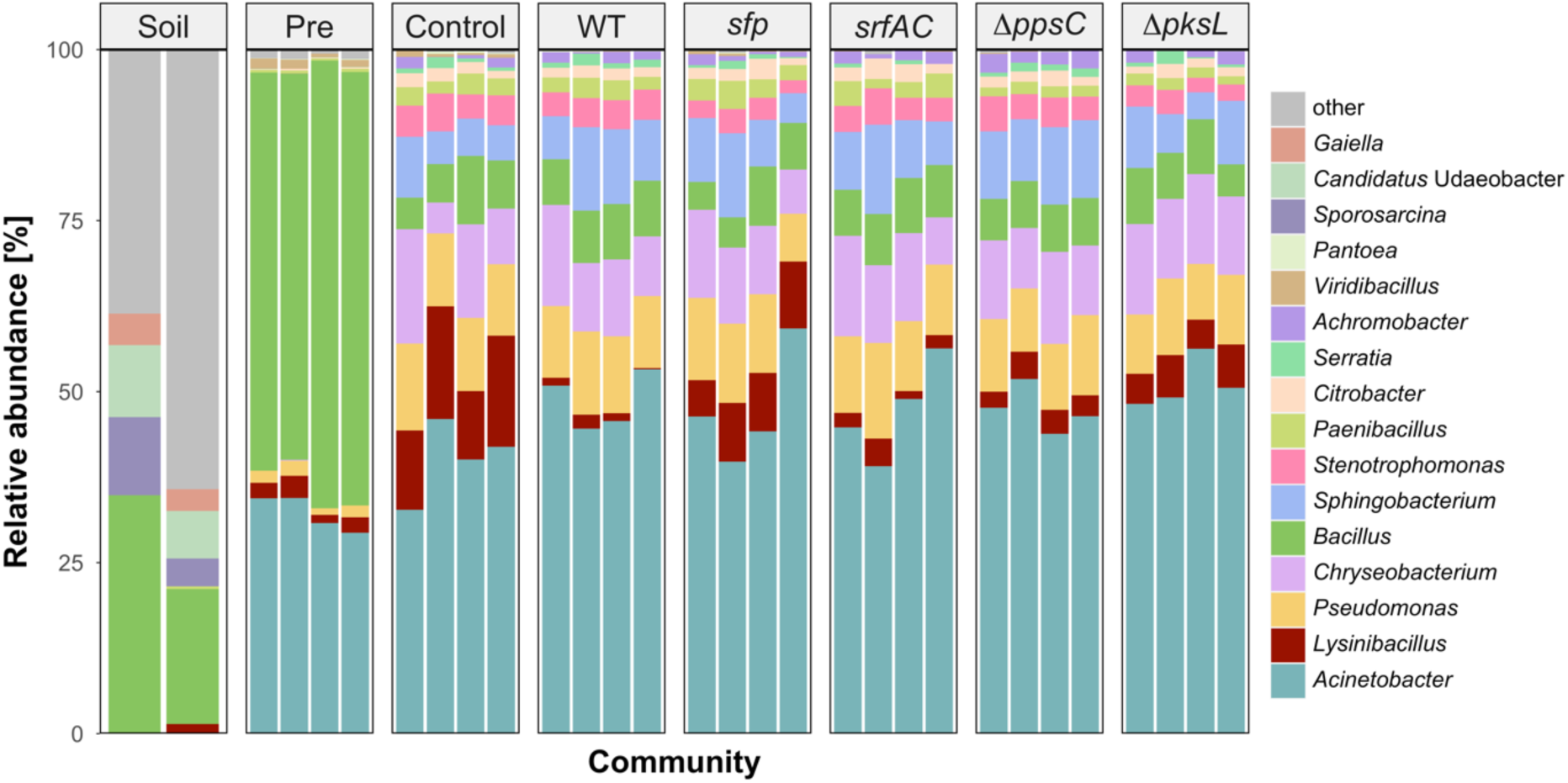
Taxonomic summaries are showing the relative abundance of the most abundant genera for each replicate of the soil sample (Soil), 12 h pre-incubated soil suspensions (Pre), and untreated mock communities (Control) or treated mock communities with either *B. subtilis* wild type (WT), NRPs deficient strain (*sfp*), surfactin mutant (*srfAC*), plipastatin mutant (Δ*ppsC*) or bacillaene mutant (Δ*pksL*) co-cultivated for 48 h. Genera are classified as “other” when the relative abundance is < 2 % (Soil), < 1 % (Pre) or < 0.19 % (in all differently treated mock communities).

Diversity analyses were performed to determine the overall impact of NRPs on the diversity of the bacterial mock communities. The read numbers varied among the different samples (Table S3), but we had to exclude the sample “Soil 1” from the analysis since it had the lowest read number and its rarefaction curve was not reaching a clear asymptote (Figure S1). The alpha diversity revealed that the mock communities co-cultivated for 48 h had Shannon indexes between 2.7 and 3.3, thus a similar genus richness and evenness (Figure 3A). However, it also highlighted that the pre-cultivated communities had the lowest Shannon indexes between 1.8 and 2.1. Consequently, these communities have a lower species evenness and are therefore dominated by fewer species. The soil sample had the highest Shannon index (6.3), which highlights that richness and evenness are expectedly larger than in the *in vitro* communities. The alpha diversity of the soil sample and pre-incubated soil suspensions differed from the mock communities, but we could not see differences among the mock communities.

**Figure 3.**
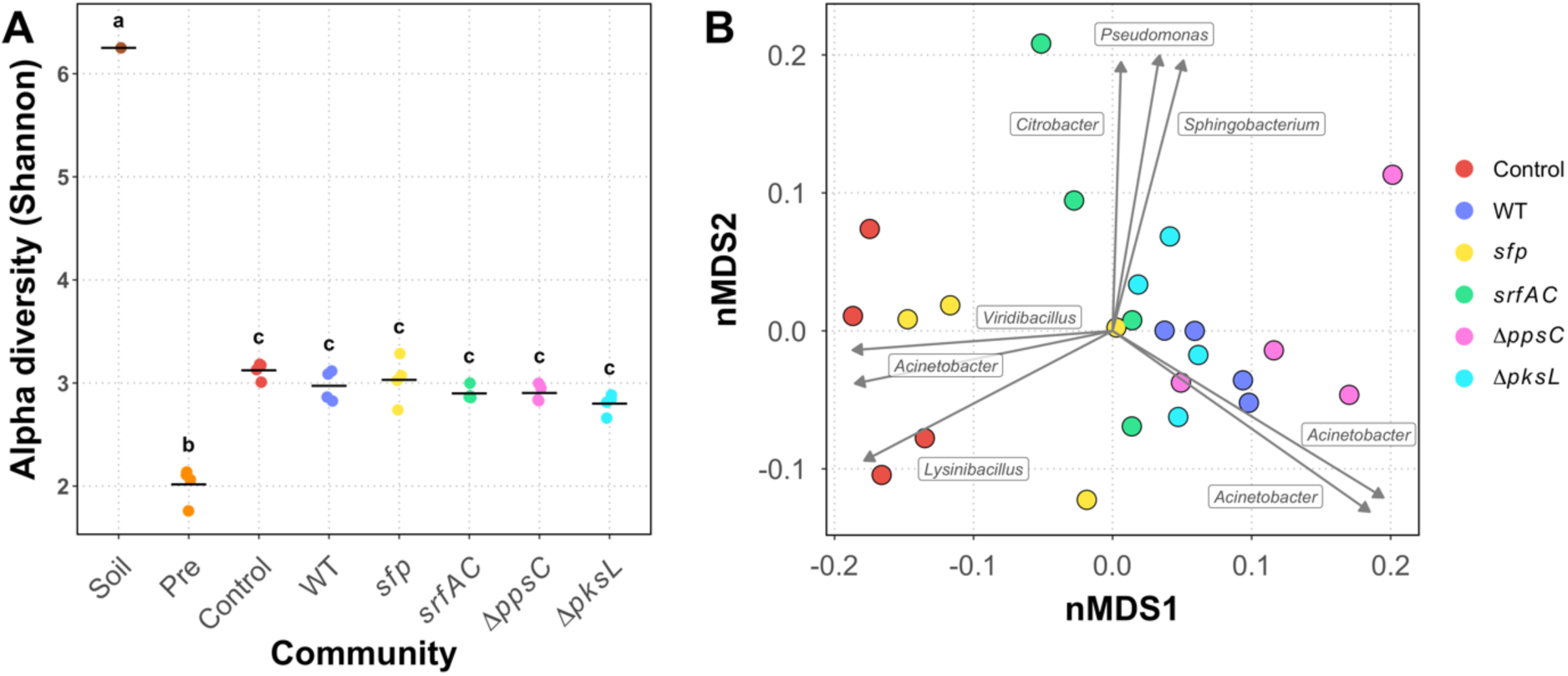
Diversity analyses of the soil sample (Soil), 12 h pre-incubated soil suspensions (Pre), and untreated mock communities (Control) or treated mock communities with either *B. subtilis* wild type (WT), NRPs deficient strain (*sfp*), surfactin mutant (*srfAC*), plipastatin mutant (Δ*ppsC*) or bacillaene mutant (Δ*pksL*) co-cultivated for 48 h. (A) Alpha diversity (Shannon) of the different samples, denoted by the x-axis. Each point represents a replicate while the line indicates the mean of the Shannon diversity indexes. (B) Beta diversity of the mock communities calculated with the Bray–Curtis dissimilarity and visualised as circles in a nonmetric multidimensional scaling (nMDS). The vectors, each labelled with the corresponding genus, represent the ASVs with the highest correlating with the nMDS ordination. Vector lengths are proportional to the level of correlation.

Therefore, we determined the beta diversity only for the treated and untreated mock communities co-cultivated for 48 h. The analysis underlined a high similarity in the composition of the mock communities treated with *B subtilis* strains (Figure 3B). However, the control mock communities separated from the majority of the treated communities along the nMDS1 axis. Interestingly, two replicates of the *sfp*-treated communities had a low Bray-Curtis dissimilarity to the control communities, emphasising a high similarity to the untreated control communities. In contrast, the communities supplemented with NRPs-producing *B. subtilis* strains clustered together and indicated a lower dissimilarity to each other than to the control communities. Notably, the communities treated with the *srfAC* mutant had a higher dispersion, likely owing to a low number of reads in two of the replicates. We fitted the most correlating (R^2^ > 0.6) amplicon sequence variants (ASVs) to the nMDS ordination and plotted them as vectors to investigate differences between the mock communities. The analysis indicated that three ASVs, taxonomically assigned to the genera *Lysinibacillus, Acinetobacter* and *Viridibacillus* correlated with the control and two *sfp*-treated communities. This observation suggests that the absence of NRPs-producing *B. subtilis* resulted in an increased abundance of these. Furthermore, two ASVs of the genus *Acinetobacter* correlated best with the communities supplemented with the NRPs-producing *B. subtilis* strains, hinting a higher frequency of them in NRPs-treated communities. Additionally, three ASVs identified as *Pseudomonas, Citrobacter* and *Sphingobacterium* correlated with two communities treated with the surfactin mutant. A similar but smaller correlation with two bacillaene mutant-treated communities was detectable as well. These results imply a negative impact of either surfactin or bacillaene on the four ASVs. Interestingly, the vector-based analysis suggests that, depending on the ASVs, the genus *Acinetobacter* is both positively and negatively affected by the NRPs.

In conclusion, the alpha diversity analyses revealed that species richness and evenness were reduced in the *in vitro* communities compared to the soil community. Furthermore, 12 h pre-incubated soil suspensions showed a reduced diversity compared to the mock communities incubated for 48 h. Nevertheless, we could not detect an effect of the supplemented *B. subtilis* strains on diversity. However, the beta diversity results suggested that the addition of NRPs-producing *B. subtilis* strains influenced the composition of the mock communities. Mainly ASVs belonging to the genera *Lysinibacillus, Viridibacillus* and *Acinetobacter* were affected by the presence or absence of *B. subtilis* NRPs in the bacterial mock communities.

The diversity, in particular the species evenness, increased in all established mock communities independent of the treatment compared to the pre-cultivated soil suspensions and contained 11 - 18 genera (Figure 2). The most abundant genera having a proportion greater than 0.19 % in at least one *B. subtilis*-treated or untreated mock community were *Acinetobacter, Lysinibacillus, Pseudomonas, Chryseobacterium, Bacillus, Sphingobacterium, Stenotrophomonas, Paenibacillus, Citrobacter, Serratia, Achromobacter, Viridibacillus* and *Pantoea*. Noteworthily, the prevalence of the *Bacillus* genus was comparable in the *B. subtilis*-treated communities (4 – 9 %) and the control (5 – 10 %). In the latter, the present *Bacillus* ssp. originated only from the soil suspension, highlighting that the additional supplementation of *B. subtilis* did not affect the relative abundance of the genus *Bacillus* after 48 h co-cultivation. Interestingly, the only genera detected in both the *in vitro* mock communities and the soil samples were *Bacillus, Lysinibacillus* and *Paenibacillus.* The remaining most abundant genera in the mock communities were below the detection limit. Comparisons of abundance ratios between the control communities and *B. subtilis* WT-treated communities revealed that *Lysinibacillus* and *Viridibacillus* were significantly decreased 9.4 (P ≤ 0.001) and 8.3-fold (P ≤ 0.01), respectively, in the communities supplemented with *B. subtilis* WT (Figure 4A). None of the other genera was affected by the addition of this strain. In comparison, we could only detect a 1.8-fold significant reduction (P ≤ 0.05) of *Lysinibacillus* in the *sfp*-treated communities compared to the untreated communities, thus greatly diminished effect compared to the WT treated samples (Figure 4B). Also, we could not observe a significant reduction of *Viridibacillus*, but besides *Lysinibacillus* also *Stenotrophomonas* was 1.7-fold (P ≤ 0.05) significantly reduced in these communities. The direct comparison of WT- and *sfp*-treated communities confirmed the NRPs-dependent suppression of both *Lysinibacillus* and *Viridibacillus* in the WT-treated communities and the suppression of *Stenotrophomonas* in the *sfp*-treated communities (Supporting Information File 2, Figure S2).

**Figure 4.**
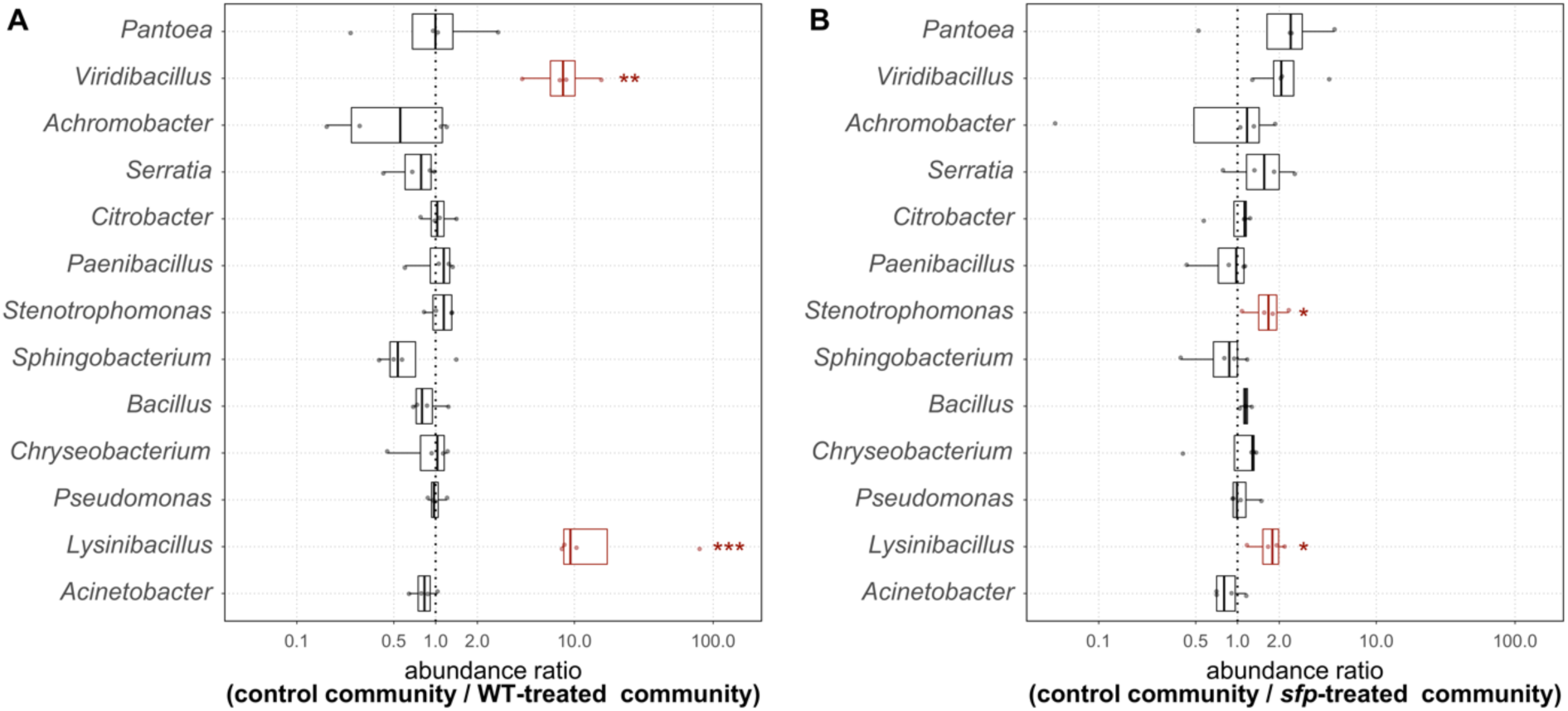
Abundance ratios for each genus and replicate (points) in the (A) control community compared to the WT-treated community and (B) control community compared to the *sfp*-treated community. Red box plots highlight statistical significance, which is defined as P ≤ 0.05 (*), P ≤ 0.01 (**) and P ≤ 0.001 (***).

Concentrating on *Lysinibacillus*, the highest abundance of this genus was discernible in the control assays (13.9 %), which was significantly different compared to all other *B. subtilis*-treated assays (Figure 5). However, when *B. subtilis* P5_B1 WT was added to the mock communities, a significant decrease (P ≤ 0.001) of *Lysinibacillus* (1.2 %) was discovered compared to the control communities. Furthermore, when we added the NRPs deficient strain *sfp*, we could notice a significantly higher abundance of *Lysinibacillus* (8.6 %) compared to the WT-treated communities (P ≤ 0.001), but still, a significantly lower prevalence compared to the control communities (P ≤ 0.05). The frequency of *Lysinibacillus* was slightly but not significantly higher in the communities treated with the single NRP mutants *srfAC* (2.0 %) and Δ*ppsC* (3.3 %) compared to the WT-treated communities. *Lysinibacillus’* abundance in the assays containing the Δ*pksL* strain (5.3 %) was significantly higher (P ≤ 0.01) than in the WT-treated assays. However, its abundance in Δ*pksL*-treated communities was not significantly different from the Δ*ppsC* or *sfp*-treated communities. In summary, *Lysinibacillus* was affected by the addition of *B. subtilis* independent of the NRPs, but when *B. subtilis* strains capable of producing them were present, the impact on *Lysinibacillus* was enhanced. Furthermore, the results indicate that bacillaene had the strongest and surfactin the weakest effect on *Lysinibacillus* in the mock communities.

**Figure 5.**
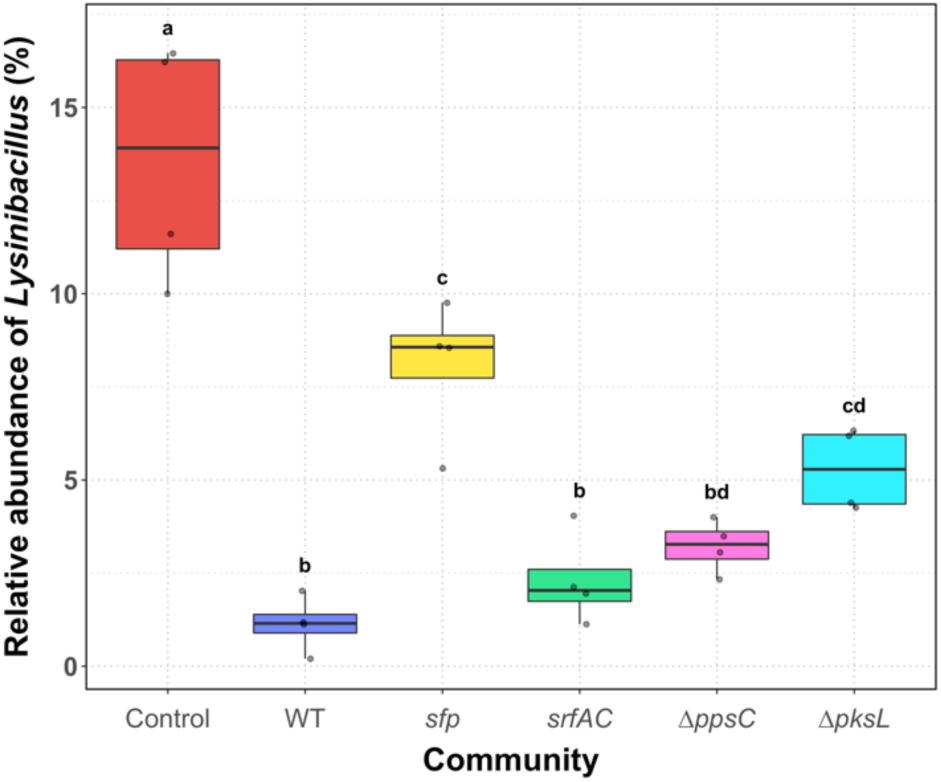
The relative abundance of *Lysinibacillus* in the untreated mock communities (Control) and treated mock communities with either *B. subtilis* wild type (WT), NRPs deficient strain (*sfp*), surfactin mutant (*srfAC*), plipastatin mutant (Δ*ppsC*) or bacillaene mutant (Δ*pksL*) co-cultivated for 48 h. Points represent the abundance in each replicate. Treatments with different letters are significantly different (P ≤ 0.05).

The second genus affected by the addition of *B. subtilis* was *Viridibacillus*, which had compared to *Lysinibacillus* a very low abundance in the control mock communities (0.49 %) (Supporting Information File 3, Figure S3). However, when *B. subtilis* WT was added to the community, *Viridibacillus* indicated a significantly lower (P ≤ 0.01) abundance (0.03 %) compared to the control communities. Notably, in two of the WT-treated community replicates, *Viridibacillus* was below the detection level. Nevertheless, the abundance of this genus in the *sfp*-treated communities (0.26 %) was statistically not significant in comparison to the WT and the control communities. Furthermore, the addition of the single NRP mutants *srfAC*, Δ*ppsC* and Δ*pksL* resulted in communities with *Viridibacillus* frequencies similar to the WT-treated communities (0.08%, 0.05% and 0.00 %, respectively). *Viridibacillus* such as *Lysinibacillus* was affected by the addition of *B. subtilis* to the communities. However, no particular NRP could be assigned to the reduced frequency of *Viridibacillus*.

### Growth properties of *L. fusiformis* M5 supplemented with *B. subtilis* spent media

The main finding from the semi-synthetic mock community experiment indicated that the genus *Lysinibacillus* was negatively affected by the addition of *B. subtilis* P5_B1 WT and that NRPs enhance the suppression. To dissect the direct impact of particular NRP in this inhibition, we monitored the growth of *L. fusiformis* M5, a previously isolated *Lysinibacillus* species [54], over 24 h treated with different proportions of spent media from *B. subtilis* WT and its NRPs mutants (Figure 6). When we added 52.80 % spent medium to *L. fusiformis,* we observed the fastest entry into the exponential phase growth in the untreated assay. Interestingly, the addition of spent medium of either WT, Δ*ppsC* or Δ*pksL* caused in *L. fusiformis* a delay into this growth phase by approximately 11-13 h compared to the control. Such a strong effect was not observed when the spent medium of the *sfp,* or *srfAC* mutant was added. The addition of these two spent media caused only a slight delay of the exponential phase growth of *L. fusiformis*, although *sfp* spent medium had a lower effect on *L. fusiformis* compared to *srfAC* spent medium. When 23.00 % spent medium was added, no growth differences could be detected anymore between the control and *sfp*-treated assays in the exponential growth phase. Furthermore, the effect of WT spent medium seems to be reduced at this concentration, but the spent media of Δ*ppsC* and Δ*pksL* maintained their growth inhibition potential. The lowest concentration of spent media having an inhibitory effect was 10.02 %. At this concentration, only the spent media of Δ*ppsC* and Δ*pksL* affected the growth of *L. fusiformis* even though it was weakened compared to higher concentrations. Intriguingly, a higher level of aggregation was observed in the *L. fusiformis* assays supplemented with the spent medium of *sfp* compared to the other assays, which caused higher and variable OD measurements in the stationary phase of the growth curves (Supporting Information File 4, Figure S4). Finally, it was noted that the final cell densities were slightly higher in the assays supplemented with the spent medium than in the control assays.

**Figure 6.**
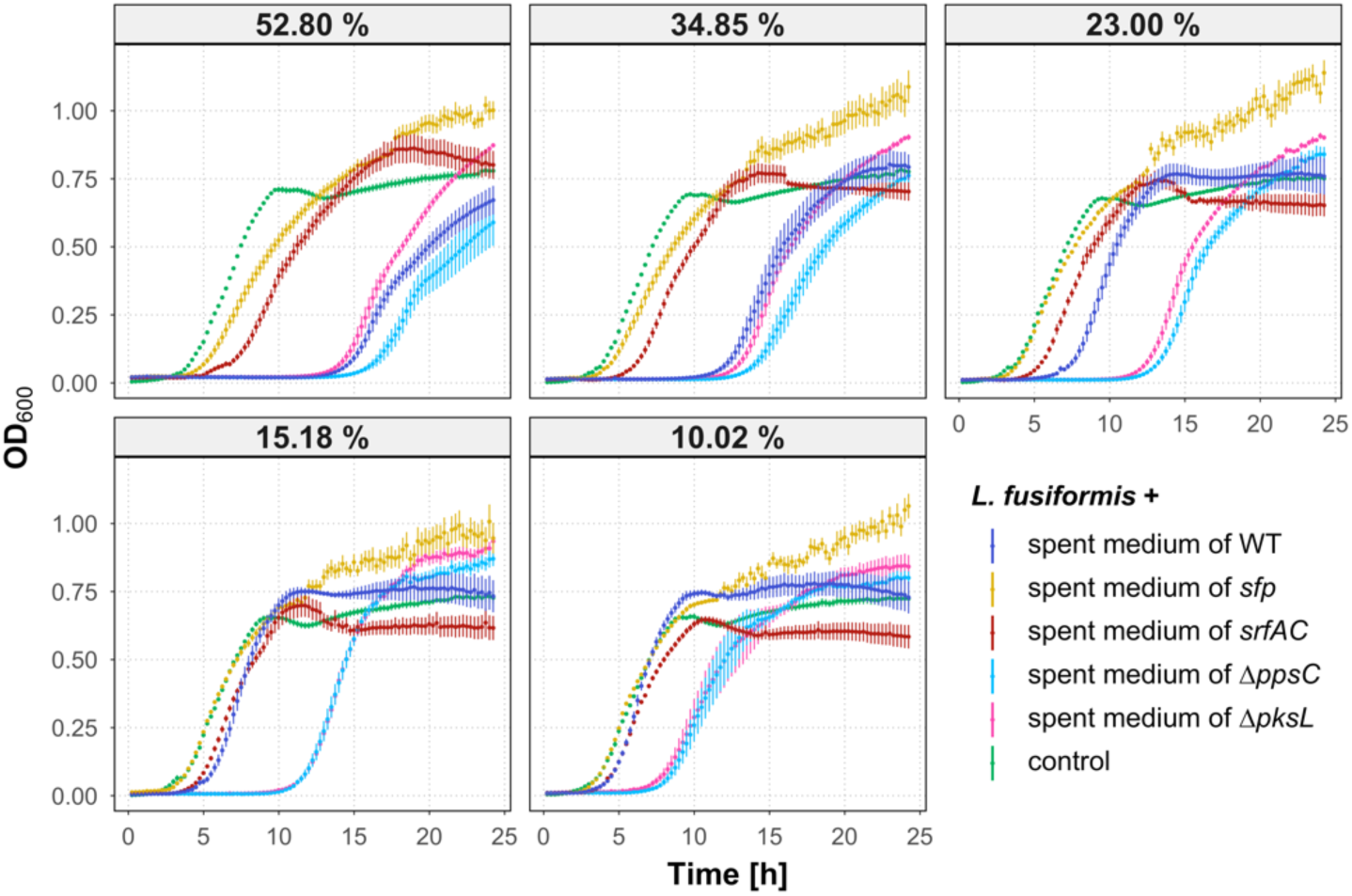
Growth curves of *L. fusiformis* M5 exposed to spent media from 48 h *B. subtilis* cultures and without treatment (control). The spent media concentrations 10.02 % to 52.80 %, acquired with a serial dilution, are indicating the proportion of spent medium from the total volume. Error bars represent the standard error. N ≥ 6.

These results revealed that *B. subtilis*-mediated inhibition of *L. fusiformis* is NRPs dependent since the spent medium of the NRPs deficient strain *sfp* had an only minor impact. Moreover, we hypothesise that surfactin is responsible for the direct inhibitory effect on *L. fusiformis*, as this was the only spent medium of an NRP mutant strain with lowered inhibition compared to spent media of other single NRP mutants.

### Impact of surfactin on the growth of *L. fusiformis*

To confirm the inhibitory effect of surfactin on *L. fusiformis*, we exposed this strain to different concentrations of pure surfactin dissolved in methanol and monitored its growth over 24 h. Growth of *L. fusiformis* was delayed in the exponential growth phase when surfactin was supplemented in concentrations between 31.25 and 500 µg/mL (Figure 7). At a concentration of 500 µg/mL surfactin, the cell density in the stationary phase was lower than the control. At a concentration of 250 µg/mL, the cell density reached a level similar to the untreated control. However, when surfactin was added in concentrations between 125 and 31.25 µg/mL, after an initial growth delay in the exponential phase, the cell densities in all treatments exceeded the ones of the control. The highest concentration of the solvent methanol (5 %) had only a minor inhibiting effect on *L. fusiformis,* whereas lower concentrations of methanol showed no inhibition (Supporting Information File 5, Figure 5). These results suggest that surfactin has growth inhibitory effects on *L. fusiformis,* and we hypothesise that it might act as the key inhibitory *B. subtilis* NRP under the tested conditions.

**Figure 7.**
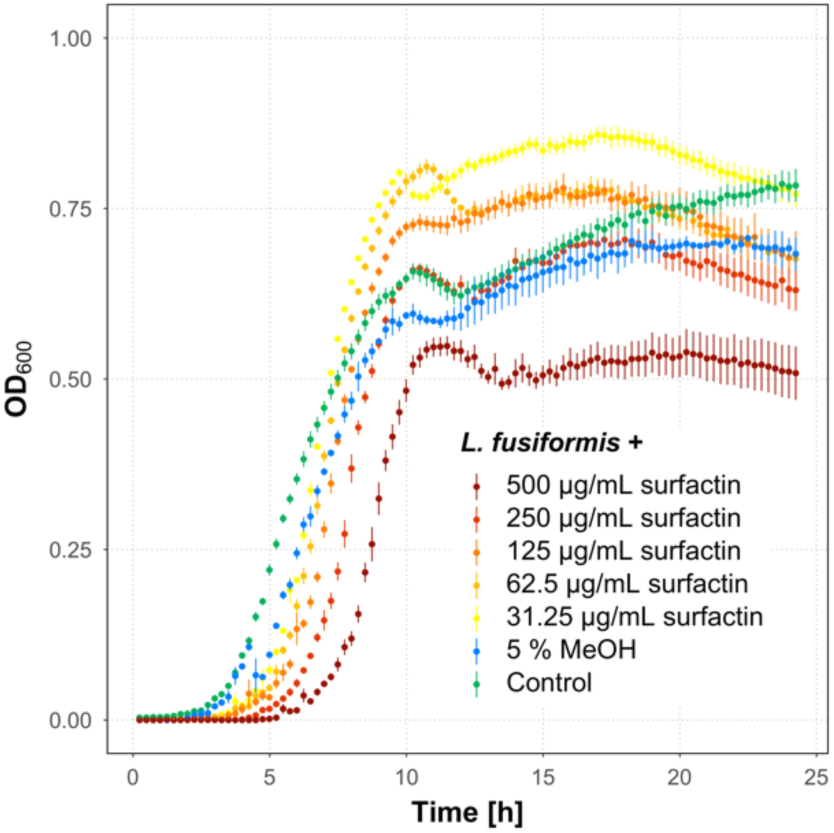
Growth curves of *L. fusiformis* M5 exposed to different concentrations of surfactin, the highest concentration of the solvent MeOH and without treatment (control). Error bars represent the standard error. N ≥ 5 (control and surfactin-treated assays), N = 2 (MeOH-treated assays).

## Discussion

*B. subtilis* is known to produce a wide range of different SMs, which target a large number of various micro and macroorganisms [35]. Our study demonstrates that the NRPs produced by the recently isolated environmental strain of *B. subtilis* P5_B1 did not strongly impact the overall soil-derived semi-synthetic mock community but reduced the abundance of the genera *Lysinibacillus* and *Viridibacillus*. Moreover, it reveals that the strain *L. fusiformis* M5 was directly affected by the *B. subtilis* lipopeptide surfactin in a monitored growth experiment.

We studied bacterial community compositions by sequencing the two variable regions V3 and V4 of the 16S rRNA gene. Noteworthy, some limitations of this technique are well-known. Poretsky *et al.* (2014) revealed that amplicon sequencing of the 16S rRNA gene indicates a lower sequence diversity and substantial differences in relative abundances of specific genus-assigned taxa compared to metagenomics. Moreover, 16S amplicon sequencing of single variable regions rarely allow sufficient discrimination below the family or genus level and therefore, intra-genus differentiation and heterogeneity cannot be addressed [55]. Furthermore, the fundamental problem is that bacteria harbour various copy numbers of the 16S rRNA gene in their genomes, which biases quantification studies [56]. Alpha diversity analyses based on the Shannon estimation revealed that diversity was strongly reduced in *in vitro* cultivations. Furthermore, it disclosed that the pre-cultured soil suspension had the lowest diversity index, because mainly the genera *Bacillus* and *Acinetobacter* were enriched, which can be probably traced back to different growth rates among the present species. A substantial shift in the community compositions was observed between *in vivo* and *in vitro* communities since the majority of genera present in the *in vitro* communities were below the detection limit in the soil sample. However, during the 12h-pre-cultivation of the soil suspension, bacteria were exposed to different nutrient availability, changed physical conditions such as temperature, liquid environment and loss of the spatial soil structure. These conditions were most likely selecting for generalist bacteria capable of proliferating under the given conditions and independent of other bacteria. During the following 48h co-cultivation, depletion of the primary nutrient sources and metabolic cross-feeding further shaped the community assembly. Goldford *et al.* (2018) revealed that the main sources of metabolic cross-feeding are secreted metabolic by-products from the community members. They further highlighted that bacterial communities stabilised after approximately eight to nine 48 h-co-cultivations [57]. In our study, bacterial communities were only co-cultivated once for 48 h, suggesting that the assembly of the bacterial communities has not yet reached a stable phase, which explains the differences between the pre-cultures and co-cultivated mock communities.

The Shannon index showed no differences among the established and differently treated mock communities, which primarily consisted of 13 genera. Even though *Bacillus* was the most abundant genus in the pre-cultures, further incubation for 48 h resulted in decreased relative abundance independently if the respective *B. subtilis* strains were seeded or the pre-cultures were untreated. It shows that the initial dominance of *Bacillus* could not be maintained at prolonged incubation. The *B. subtilis* strains were added at a community assembly phase when *Bacillus* was the dominating genus so that the general genera distribution was not expected to be influenced extensively. Nevertheless, after 48 h co-cultivation, the final relative abundance of the *Bacillus* genus was not increased in the communities treated with *B. subtilis* compared to the control. This observation highlights that the presence or absence of NRPs did not affect the competitiveness of *B. subtilis*. However, the 16S amplicon sequencing did not allow the detection of interactions and competitions within the *Bacillus* genus. The composition of this genus could vary among the differently treated communities. Nonetheless, the beta diversity analysis indicated a dissimilarity between the untreated and treated mock communities. Besides, two of the communities treated with the *sfp*-mutant showed the highest similarity to the untreated communities suggesting that the supplementation of the NRPs-producing *B. subtilis* strains affected the communities. The vectors of *Acinetobacter* ASVs had a direction either to NRPs-treated or NRPs-untreated communities, indicating that the NRPs influenced species within the same genus differently.

In microbial communities, the amount of interactions and relations increases with the number of community members. The established semi-synthetic mock communities in this study contained at least 13 genera with a relative abundance > 0.19 %. Therefore, it can be assumed that various interactions between them occurred. Nevertheless, we could observe statistically significant reductions of the two genera, *Lysinibacillus* and *Viridibacillus*, in communities supplemented with the NRPs-producing *B subtilis* wild type strain. In contrast, in communities supplemented with the NRPs deficient mutant *sfp, Lysinibacillus* was more frequent than in the wild type-treated communities. This observation indicates that NRPs have a great impact on suppressing *Lysinibacillus*. However, further factors are involved in the suppression as well, since the *sfp* mutant maintained a reduction of *Lysinibacillus*, even though weakened. Moreover, no particular NRP could be allocated to the inhibition of the *Lysinibacillus* genus in these semi-synthetic communities, but bacillaene displayed the highest impact on the suppression. An inhibition of *Viridibacillus* mediated by NRPs was also observable, but for this genus, bacillaene had the lowest impact. However, these results must be interpreted with caution and need further investigations, since *Viridibacillus* was one of the lowest abundant genera in the mock communities and abundance calculations are sensitive to the depth of sequencing. Besides the suppression of *Lysinibacillus* and *Viridibacillus, Stenotrophomonas* was uniquely suppressed in the communities supplemented with the *sfp* mutant but not when the WT strain was added. This observation might be evoked by inhibiting other species which, in turn, has lessened inhibition of *Stenotrophomonas*.

Previous studies revealed that the introduction of SM-producing bacteria to a bacterial community had no major impact on the entire composition. The marine bacterium *Phaeobacter inhibens*, producing tropodithietic acid (TDA), did not strongly influence the microbiome diversity of the oyster *Ostrea edulis* but reduced the relative abundance of the orders Vibrionales and Mycoplasmatales [58]. Similar results were achieved when *B. velezensis* FZB42 was successfully applied as a biocontrol agent to lettuce in soil [59]. The authors could not see a substantial impact on the rhizosphere bacterial community by the supplemented biocontrol strain, whereas sampling time and additional inoculation of the fungal plant pathogen influenced the community to a greater extent. Apart from soluble SM, volatile organic compounds (VOCs) are as well capable of impacting a microbial community. Cosetta *et al.* (2020) demonstrated that VOCs of cheese rind-associated fungi have both growth stimulating and inhibiting properties on members of the rind microbiome. The authors could reveal that the VOC-mediated shift of the bacterial community was caused due to growth promotion of *Vibrio* spp. [60]. These studies and the results from the semi-synthetic mock community experiment of this study highlight that the overall impact of SMs on the targeted microbial communities is low, which suggest being no mass destruction compounds. However, in all communities, distinct genera or species were suppressed or promoted, emphasising the potential of SMs to shape microbial communities.

To investigate if *Lysinibacillus* is sensitive to any particular NRP of *B. subtilis*, we exposed the isolate *L. fusiformis* M5 to the spent media of the respective *B. subtilis* strains and monitored the growth. *L. fusiformis* M5 has been isolated from soil and demonstrated to impact biofilm colony development of *B. subtilis* [54]. Interestingly, the modulation of biofilm development was mediated by the primary metabolite hypoxanthine secreted by *L. fusiformis*. Of note, the impact of *B. subtilis* was not noticed on *L. fusiformis* in the mixed colony biofilm communities, possibly due to the use of an NRPs negative *B. subtilis* strain, 168 that harbours a spontaneous frameshift mutation in *sfp* gene [54]. Testing the impact of the natural isolate *B. subtilis* P5_B1 and its NRP mutant derivatives revealed that the spent media from both the NRPs deficient strain *sfp* and the surfactin deficient strain *srfAC* had the lowest impact on the growth of *L. fusiformis*. In addition, the spent media of Δ*ppsC* and Δ*pksL* maintained their bioactivity at low concentrations, whereas the effect of WT was already strongly reduced at this level of the spent medium. This difference could occur, on the one hand, due to higher levels of surfactin in the two mutants compared to the WT. On the other hand, the spent medium originated from cultures with an optical density at 600 nm (OD_600_) of 3.0. Cultures with higher ODs were diluted before the harvesting and WT cultures exhibited overall, the highest ODs among the strains. Since the NRPs concentration is not proportional to the final OD due to, e.g. occurring of cell lysis, the spent media might be slightly differently diluted among the strains. Therefore, minor differences might be observable in the assays supplemented with highly diluted spent media. The observation that *L. fusiformis* displays slightly higher cell density when the bacterial spent medium is supplemented might be due to the availability of additional nutrients. Nevertheless, the supernatant and pure compound supplementation demonstrated that surfactin is a direct suppressor of *L. fusiformis*. However, as the spent media of the *sfp* and *srfAC* strains still had growth inhibition effect, it is plausible that next to surfactin further NRPs and even other compounds might provoke slight growth suppression of *Lysinibacillus*. When *L. fusiformis* was exposed to surfactin concentrations between 31.25 and 125 µg/mL, higher final cell densities were detectable than in assays treated with higher levels of surfactin or in the control. Interestingly, Arjes *et al.* (2020) recently demonstrated that surfactin enhances the availability of oxygen for *B. subtilis* by increasing oxygen diffusivity [61], which might also positively affect the growth of *L. fusiformis*.

Experiments with differently treated semi-synthetic mock communities have demonstrated that *Lysinibacillus* and *Viridibacillus* were affected by the addition of an NRPs producing *B. subtilis* strain. *Lysinibacillus* was least affected in the mock communities supplemented with the *B. subtilis* Δ*pksL* strain incapable of producing bacillaene, suggesting that bacillaene is the most active compound against this genus. In contrast, the growth curve experiments showed that *L. fusiformis* M5 is most sensitive to surfactin. Importantly, our analysis does not reveal which *Lysinibacillus* species were present in the mock communities, and therefore their sensitivity might be different from the test species used, *L. fusiformis*. Moreover, the spent medium was harvested from pure cultures of *B. subtilis* grown in an undiluted complex medium which might change the production of NRPs due to lacking impact of the community members and the level of nutrients. Thus, lower concentrations of the NRPs in the mock communities might affect *Lysinibacillus* differently compared to the monoculture growth experiments supplemented with spent media. Finally, *Lysinibacillus* can also be affected indirectly by *B. subtilis* NRPs in the mock communities. Bacillaene is described as a wide-spectrum antibiotic disrupting the protein synthesis in bacteria [34],[52]. The observations suggest that it has the most substantial impact on specific members of the mock community and consequently, an indirect effect on *Lysinibacillus*. Nevertheless, the exact mechanisms at play remain to be deciphered.

Interestingly, the two genera, *Lysinibacillus* and *Viridibacillus*, of the mock communities are, besides *Paenibacillus*, the closest relatives of *B. subtilis*. The fact that suppression effects are only observable for these genera could presumably be caused by the higher overlap in the ecological niches, triggering competition for the same nutrients. Indeed, a higher phylogenetic and metabolic similarity between bacteria increases the probability of antagonism [62].

We could not quantify the concentrations of *B. subtilis* NRPs in the mock communities, since the detection of low concentrations is still under development. However, a better understanding of their impact on the mock communities could be realised by further experiments investigating the effect of supplemented pure NRP compounds, e.g. surfactin and bacillaene. The impact of antibiotics on algae-associated bacterial communities was investigated by Geng *et al.* (2016), who revealed a dose-depended influence of pure TDA on the microbiome structure of *Nannochloropsis salina* [63]. Such pure NRP supplementations in various concentrations would allow exploring their effects on bacterial community assembly. Furthermore, *in vivo* experiments could reveal the impact of NRPs on microbial communities in complex natural systems similar to the study from Chowdhury *et al.* (2013) [59]. Noteworthy, our study focused only on NRPs, but additional SMs, such as bacteriocins, are as well predicted for *B. subtilis* P5_B1 [51]. Future investigations should investigate the impact of both bacteriocins and NRPs on microbial communities.

## Conclusion

In summary, this study demonstrates that nonribosomal peptides of *B. subtilis* P5_B1 have only minor impact on the overall structure of soil-derived semi-synthetic bacterial mock communities but suppress significantly the genera *Lysinibacillus* and *Viridibacillus*. Furthermore, it highlights the bioactivity of surfactin against *L. fusiformis* M5.

## Experimental

### Strains, media and chemicals

All strains used in this study are listed in Table S1 (Supporting Information File 1). For routine growth, bacterial cells were cultured in tryptic soy broth (TSB, CASO Broth, Sigma-Aldrich) containing 17 g l^−1^ casein peptone, 3 g l^−1^ soy peptone, 5 g l^−1^ sodium chloride, 2.5 g l^−1^ dipotassium hydrogen phosphate and 2.5 g l^−1^ glucose.

### Semi-synthetic mock community assay

Semi-synthetic soil communities were obtained from the soil of sampling site P5 (55.788800, 12.558300) [51],[64]. 1 g soil was mixed in a 1:9 ratio with 0.9 % saline solution, vortexed on a rotary shaker for 15 min and allowed to sediment for 2 min. Four independent communities were established by inoculating 10-times diluted TSB (0.1x TSB) with 1 % soil suspension taken from the middle part of the liquid phase, followed by incubation at 21-23°C and 250 rpm for 12 h. Simultaneously, pre-grown *B. subtilis* P5_B1 WT and its NRP mutant derivatives were inoculated in 0.1x TSB and incubated parallel using the same conditions. After 12 h pre-cultivation, 3 mL aliquots of the soil suspension were transferred into six glass tubes. One tube was left untreated and functioned as control, whereas the remaining five were supplemented with respective *B. subtilis* strains by adding 10 % of the final volume. The cultures were incubated at 21-23°C and 250 rpm for 48 h. DNA was extracted from two replicates of the initial soil sample, the 12-h-pre-cultivated soil suspensions and the *B*. subtilis-treated or untreated mock communities co-cultivated for 48 h.

### DNA extraction

Environmental and semi-synthetic community genomic DNA was extracted from either 250 mg soil or 250 µl bacterial culture, respectively by using the DNeasy PowerSoil Pro Kit (QIAGEN) and following the manufacturer’s instructions.

### Amplification of 16S rRNA hypervariable regions V3-V4

The V3-V4 region of the 16S rRNA gene was PCR amplified from the extracted DNA samples using Fw_V3V4 (5’-CCTACGGGNGGCWGCAG-3’) and Rv_V3V4 (5’-GACTACHVGGGTATCTAATCC-3’) primers that were tagged with short barcodes with a length of eight nucleotides, listed in Table S2 (Supporting Information File 1). The PCR reactions contained 10.6 μl DNase free water, 12.5 μl TEMPase Hot Start 2x Master Mix, 0.8 μl of each primer (10 μM) and 0.3 μl of 50 ng/µl DNA template. The PCR was performed using the conditions of 95°C for 15 min followed by 30 cycles of 95°C for 30 s, 62°C for 30 s, 72°C for 30 s, and finally, 72°C for 5 min. All V3-V4 amplicons were purified using the NucleoSpin Gel and PCR Clean-up kit (Macherey-Nagel) and pooled in equimolar ratios. The amplicon pool was submitted to Novogene Europe Company Limited (United Kingdom) for high-throughput sequencing on an Illumina NovaSeq 6000 platform with 2 million reads (2x 250 bp paired-end reads). Raw sequence data is available at NCBI: PRJNA658074.

### Sequencing data pre-processing

The multiplexed sequencing data was imported into the QIIME 2 pipeline (version 2020.6) [65],[66]. The paired-end sequences were demultiplexed with the QIIME 2 plugin cutadapt [67]. The minimum overlap of partial matches between the read and the barcode sequence was set to 5 nucleotides to reduce random matches. The QIIME 2 implementation DADA2 was used to denoise and merge paired-end reads [68]. In total, 362,475 reads were assigned to the respective samples with an average of 12,083 reads per sample (range: 751 to 34,802; Supporting Information File 1, Table S3. The 16S rRNA reference sequences with a 99 % identity criterion obtained from the SILVA database release 132 were trimmed to the V3-V4 region, bound by the primer pair used for amplification and the product length was limited to 200-500 nucleotides [69]. Taxonomy was assigned to the sequences in the feature table generated by DADA2 by using the VSEARCH-based consensus taxonomy classifier [70]. A tree for phylogenetic diversity analyses was generated with FastTree 2 from the representative-sequences [71]–[73].

### Relative species abundance and phylogenetic diversity analyses

QIIME 2 artifacts were imported into the R software (4.0.2) with the R package qiime2R, and further analyses were conducted in the R package phyloseq [74]–[76]. The taxonomy summaries were achieved by merging ASVs of the same genera and calculating their relative abundance in each sample. Differences in the presence of the most abundant genera in the control communities, in the communities supplemented with *B. subtilis* WT as well as in the communities supplemented with *B. subtilis sfp,* were investigated by calculating the abundance ratios of the different treated communities for each replicate. If species were not detected in some of the replicates, 0 values were replaced with the lowest detected value of the genus to avoid infinite values or 0 values in the ratio calculations. Rarefaction curves of the samples were calculated and visualised with the R package ranacapa [77]. Diversity analyses of the *B. subtilis*-treated and untreated samples were performed with ASV counts multiplied with factor 100,000 and transformed into integer proportions. Alpha diversity was estimated with the Shannon diversity index in the R package phyloseq [76]. Beta-diversity was determined by dissimilarities among the samples with the Bray-Curtis distance and visualised in a nonmetric multidimensional scaling (nMDS) with the R package vegan [78]. The correlation of individual ASVs on the overall bacterial community composition was calculated with the envfit function with 999 permutations from the R package vegan. The most correlating (R^2^ > 0.6) ASVs were added to the nMDS ordination plot. All graphical visualisations were realised with ggplot2 [79].

### Statistical analysis

Statistical significance was determined with the square roots of the tested values. Normality and equality of variances were tested with Shapiro-Wilk normality test and Levene’s test, respectively. If one of the tests was rejected, the non-parametric Kruskal-Wallis rank sum test was performed instead. The statistical significance of pairs was determined with the Welch two sample t-test and the differences among groups > 2 was determined with one-way analysis of variance (ANOVA) test and Tukey’s HSD test. Statistical significance was determined with an alpha level < 0.05.

### Growth monitoring of *L. fusiformis* supplemented with *B. subtilis* spent media and pure surfactin

Spent media of *B. subtilis* strains were harvested from cultures grown in TSB medium at 37°C and 250 rpm for 48 h immediately before the growth experiments. Cultures were adjusted to OD_600_ 3.0 and centrifuged for 4 min at 5,000× g. Subsequently, the supernatants were passed through 0.22 µm filters and stored at 4°C. The growth experiments were performed in 96-well microplates. The wells of the first column were filled with 30 µl 10x TSB, 30 µl *L. fusiformis* culture adjusted to OD_600_ 0.1 in 1x TSB and 240 µl of the appropriate *B. subtilis* spent medium or water (untreated control). 100 µl *L. fusiformis* culture adjusted to OD_600_ 0.01 in 1x TSB was added to the wells of the remaining columns. A 1.5-fold serial dilution of the spent media was performed column-by-column. A surfactin stock solution was prepared by dissolving 10 mg of surfactin (Sigma-Aldrich) in 1 mL methanol (MeOH). The wells of the first column were filled with 170 µl 1x TSB, 20 µl *L. fusiformis* culture adjusted to OD_600_ 0.1 in 1x TSB, and 10 µl surfactin, 10 µl MeOH (solvent control) or 10 µl 1x TSB (untreated control). To the wells of the remaining columns, 100 µl *L. fusiformis* culture was added adjusted to OD_600_ 0.01 in 1x TSB. A 2-fold serial dilution of surfactin or MeOH was performed column-by-column. In both assays, the growth of *L. fusiformis* was monitored in a microplate reader (BioTek Synergy HTX Multi-Mode Microplate Reader). The microplates were incubated at 30°C with continuous shaking (548 cpm, 2 mm) and the OD_600_ was measured in 15 min-intervals throughout 24 h. All graphical visualisations were prepared using ggplot2 [79].

## Supporting Information

Supporting information features the bacterial strains used in this study (Table S1), the 16S rRNA V3-V4 primer list (Table S2), the number of sequencing reads per sample (Table S3) and supporting figures (Figure S1-S5).

Supporting Information File:

File Name: Supporting_information.docx File Format: docx

Title: Supporting information

## Acknowledgement

This project was supported by the Danish National Research Foundation (DNRF137) for the Center for Microbial Secondary Metabolites (CeMiSt).

The authors thank the suggestions of Lone Gram and the CeMiSt centre members on the project.

Part of the Graphical Abstract has been created using BioRender.com.

## REFERENCES

1. Falkowski, P. G., Fenchel, T.; Delong, E. F. Science (80-.). 2008, 320, 1034–1039.

2. Sunagawa, S.; Coelho, L. P.; Chaffron, S.; Kultima, J. R.; Labadie, K.; Salazar, G.; Djahanschiri, B.; Zeller, G.; Mende, D. R.; Alberti, A.; Cornejo-Castillo, F. M.; Costea, P. I.; Cruaud, C.; D’Ovidio, F.; Engelen, S.; Ferrera, I.; Gasol, J. M.; Guidi, L.; Hildebrand, F.; Kokoszka, F.; Lepoivre, C.; Lima-Mendez, G.; Poulain, J.; Poulos, B. T.; Royo-Llonch, M.; Sarmento, H.; Vieira-Silva, S.; Dimier, C.; Picheral, M.; Searson, S.; Kandels-Lewis, S.; Boss, E.; Follows, M.; Karp-Boss, L.; Krzic, U.; Reynaud, E. G.; Sardet, C.; Sieracki, M.; Velayoudon, D.; Bowler, C.; De Vargas, C.; Gorsky, G.; Grimsley, N.; Hingamp, P.; Iudicone, D.; Jaillon, O.; Not, F.; Ogata, H.; Pesant, S.; Speich, S.; Stemmann, L.; Sullivan, M. B.; Weissenbach, J.; Wincker, P.; Karsenti, E.; Raes, J.; Acinas, S. G.; Bork, P. Science (80-.). 2015, 348, 1261359.

3. Martiny, J. B. H.; Bohannan, B. J. M.; Brown, J. H.; Colwell, R. K.; Fuhrman, J. A.; Green, J. L.; Horner-Devine, M. C.; Kane, M.; Krumins, J. A.; Kuske, C. R.; Morin, P. J.; Naeem, S.; Øvreås, L.; Reysenbach, A. L.; Smith, V. H.; Staley, J. T. Nat. Rev. Microbiol. 2006, 4, 102–112.

4. Berendsen, R. L.; Pieterse, C. M. J.; Bakker, P. A. H. M. Trends Plant Sci. 2012, 17, 478–486.

5. Fuhrman, J. A. Nature 2009, 459, 193–199.

6. Friedman, J.; Higgins, L. M.; Gore, J. Nat. Ecol. Evol. 2017, 1, 109.

7. Antoniewicz, M. R. Curr. Opin. Biotechnol. 2020, 64, 230–237.

8. Flemming, H. C.; Wuertz, S. Nat. Rev. Microbiol. 2019, 17, 247–260.

9. Phillips, J. D. Soil Sci. 2017, 182, 117–127.

10. Kuzyakov, Y.; Blagodatskaya, E. Soil Biol. Biochem. 2015, 83, 184–199.

11. Whipps, J. M. J. Exp. Bot. 2001, 52, 487–511.

12. Smit, E.; Leeflang, P.; Gommans, S.; Van Den Broek, J.; Van Mil, S.; Wernars, K. Appl. Environ. Microbiol. 2001, 67, 2284–2291.

13. Dunfield, K. E.; Germida, J. J. Appl. Environ. Microbiol. 2003, 69, 7310–7318.

14. Lan, G.; Li, Y.; Lesueurd, D.; Wu, Z.; Xie, G. Sci. Total Environ. 2018, 626, 826–834.

15. Garbeva, P.; Van Elsas, J. D.; Van Veen, J. A. Plant Soil 2008, 302, 19–32.

16. Singh, B. K.; Munro, S.; Potts, J. M.; Millard, P. Appl. Soil Ecol. 2007, 36, 147–155.

17. Berg, G.; Smalla, K. FEMS Microbiol. Ecol. 2009, 68, 1–13.

18. Lundberg, D. S.; Lebeis, S. L.; Paredes, S. H.; Yourstone, S.; Gehring, J.; Malfatti, S.; Tremblay, J.; Engelbrektson, A.; Kunin, V.; Rio, T. G. Del; Edgar, R. C.; Eickhorst, T.; Ley, R. E.; Hugenholtz, P.; Tringe, S. G.; Dangl, J. L. Nature 2012, 488, 86–90.

19. Lebeis, S. L.; Paredes, S. H.; Lundberg, D. S.; Breakfield, N.; Gehring, J.; McDonald, M.; Malfatti, S.; Del Rio, T. G.; Jones, C. D.; Tringe, S. G.; Dangl, J. L. Science (80-.). 2015, 349, 860–864.

20. Bulgarelli, D.; Rott, M.; Schlaeppi, K.; Ver Loren van Themaat, E.; Ahmadinejad, N.; Assenza, F.; Rauf, P.; Huettel, B.; Reinhardt, R.; Schmelzer, E.; Peplies, J.; Gloeckner, F. O.; Amann, R.; Eickhorst, T.; Schulze-Lefert, P. Nature 2012, 488, 91–95.

21. Bai, Y.; Müller, D. B.; Srinivas, G.; Garrido-Oter, R.; Potthoff, E.; Rott, M.; Dombrowski, N.; Münch, P. C.; Spaepen, S.; Remus-Emsermann, M.; Hüttel, B.; McHardy, A. C.; Vorholt, J. A.; Schulze-Lefert, P. Nature 2015, 528, 364–369.

22. Niu, B.; Paulson, J. N.; Zheng, X.; Kolter, R. Proc. Natl. Acad. Sci. U. S. A. 2017, 114, E2450–E2459.

23. Patin, N. V; Schorn; M., Aguinaldo, K.; Lincecum, T.; Moore, B. S.; Jensen, P. R. Appl. Environ. Microbiol. 2017, 83, 2676–2692.

24. Foster, K. R.; Bell, T. Curr. Biol. 2012, 22, 1845–1850.

25. Romero, D.; Traxler, M. F.; López, D.; Kolter, R. Chem. Rev. 2011, 111, 5492–5505.

26. Linares, J. F.; Gustafsson, I.; Baquero, F.; Martinez, J. L. Proc. Natl. Acad. Sci. U. S. A. 2006, 103, 19484–19489.

27. Straight, P. D.; Willey, J. M.; Kolter, R. J. Bacteriol. 2006, 188, 4918–4925.

28. Pettit, R. K. Appl. Microbiol. Biotechnol. 2009, 83, 19–25.

29. Wakefield, J.; Hassan, H. M.; Jaspars, M.; Ebel, R.; Rateb, M. E. Front. Microbiol. 2017, 8, 1284.

30. Kovács, Á. T. Trends in Microbiology. Elsevier Ltd 2019, pp 724–725.

31. Hashem, A.; Tabassum, B.; Fathi Abd_Allah, E. Saudi J. Biol. Sci. 2019, 26, 1291–1297.

32. Gadhave, K. R.; Devlin, P. F.; Ebertz, A.; Ross, A.; Gange, A. C. Microb. Ecol. 2018, 76, 741–750.

33. Stein, T. Mol. Microbiol. 2005, 56, 845–857.

34. Kaspar, F.; Neubauer, P.; Gimpel, M. J. Nat. Prod. 2019, 82, 2038–2053.

35. Harwood, C. R.; Mouillon, J. M.; Pohl, S.; Arnau, J. FEMS Microbiol. Rev. 2018, 42, 721–738.

36. Ongena, M.; Jacques, P. Trends Microbiol. 2008, 16, 115–125.

37. Quadri, L. E. N.; Weinreb, P. H.; Lei, M.; Nakano, M. M.; Zuber, P.; Walsh, C. T. Biochemistry 1998, 37, 1585–1595.

38. Kearns, D. B.; Losick, R. Mol. Microbiol. 2003, 49, 581–590.

39. Grau, R. R.; De Oña, P.; Kunert, M.; Leñini, C.; Gallegos-Monterrosa, R.; Mhatre, E.; Vileta, D.; Donato, V.; Hölscher, T.; Boland, W.; Kuipers, O. P.; Kovács, Á. T. MBio 2015, 6, e00581–15.

40. Sheppard, J. D.; Jumarie, C.; Cooper, D. G.; Laprade, R. BBA - Biomembr. 1991, 1064, 13–23.

41. Heerklotz, H.; Wieprecht, T.; Seelig, J. J. Phys. Chem. B 2004, 108, 4909–4915.

42. Heerklotz, H.; Seelig, J. Eur. Biophys. J. 2007, 36, 305–314.

43. Loiseau, C.; Schlusselhuber, M.; Bigot, R.; Bertaux, J.; Berjeaud, J. M.; Verdon, J. Appl. Microbiol. Biotechnol. 2015, 99, 5083–5093.

44. Sabaté, D. C.; Audisio, M. C. Microbiol. Res. 2013, 168, 125–129.

45. Umezawa, H.; Aoyagi, T.; Nishikiori, T.; Yamagishi, Y.; Okuyama, A.; Hamada, M.; Takeuchi, T. J. Antibiot. (Tokyo). 1986, 39, 737–744.

46. Deleu, M.; Paquot, M.; Nylander, T. J. Colloid Interface Sci. 2005, 283, 358–365.

47. Romero, D.; De Vicente, A.; Rakotoaly, R. H.; Dufour, S. E.; Veening, J. W.; Arrebola, E.; Cazorla, F. M.; Kuipers, O. P.; Paquot, M.; Pérez-García, A. Mol. Plant-Microbe Interact. 2007, 20, 430–440.

48. Alvarez, F.; Castro, M.; Príncipe, A.; Borioli, G.; Fischer, S.; Mori, G.; Jofré, E. J. Appl. Microbiol. 2012, 112, 159–174.

49. Falardeau, J.; Wise, C.; Novitsky, L.; Avis, T. J. J. Chem. Ecol. 2013, 39, 869–878.

50. Zhang, L.; Sun, C. Appl. Environ. Microbiol. 2018, 84, e00445–18.

51. Kiesewalter, H. T.; Lozano-Andrade, C. N.; Wibowo, M.; Strube, M. L.; Maróti, G.; Snyder, D.; Jørgensen, T. S.; Larsen, T. O.; Cooper, V. S.; Weber, T.; Kovács, Á. T. bioRxiv 2020, 2020.08.05.238063.

52. Patel, P.; Huang, S.; Fisher, S.; Pirnik, D.; Aklonis, C.; Dean, L.; Meyers, E.; Fernandes, P.; Mayerl, F. J. Antibiot. (Tokyo). 1995, 48, 997–1003.

53. Müller, S.; Strack, S. N.; Hoefler, B. C.; Straight, P. D.; Kearns, D. B.; Kirby, J. R. Appl. Environ. Microbiol. 2014, 80, 5603–5610.

54. Gallegos-Monterrosa, R.; Kankel, S.; Götze, S.; Barnett, R.; Stallforth, P.; Kovács, Á. T. J. Bacteriol. 2017, 199, e00204–17.

55. Poretsky, R.; Rodriguez-R, L. M.; Luo, C.; Tsementzi, D.; Konstantinidis, K. T. PLoS One 2014, 9, e93827.

56. Louca, S.; Doebeli, M.; Parfrey, L. W. Microbiome 2018, 6, 41.

57. Goldford, J. E.; Lu, N.; Bajić, D.; Estrela, S.; Tikhonov, M.; Sanchez-Gorostiaga, A.; Segrè, D.; Mehta, P.; Sanchez, A. Science (80-.). 2018, 361, 469–474.

58. Dittmann, K. K.; Sonnenschein, E. C.; Egan, S.; Gram, L.; Bentzon-Tilia, M. Environ. Microbiol. Rep. 2019, 11, 401–413.

59. Chowdhury, S. P.; Dietel, K.; Rändler, M.; Schmid, M.; Junge, H.; Borriss, R.; Hartmann, A.; Grosch, R. PLoS One 2013, 8, e68818.

60. Cosetta, C. M.; Kfoury, N.; Robbat, A.; Wolfe, B. E. Environ. Microbiol. 2020, 1462- 2920.15223.

61. Arjes, H. A.; Vo, L.; Dunn, C. M.; Willis, L.; DeRosa, C. A.; Fraser, C. L.; Kearns, D. B.; Huang, K. C. Curr. Biol. 2020, 30, 1011-1022.e6.

62. Russel, J.; Røder, H. L.; Madsen, J. S.; Burmølle, M.; Sørensen, S. J. Proc. Natl. Acad. Sci. U. S. A. 2017, 114, 10684–10688.

63. Geng, H.; Tran-Gyamfi, M. B.; Lane, T. W.; Sale, K. L.; Yu, E. T. Front. Microbiol. 2016, 7, 1155.

64. Kiesewalter, H. T.; Lozano-Andrade, C. N.; Maróti, G.; Snyder, D.; Cooper, V. S.; Jørgensen, T. S.; Weber, T.; Kovács, Á. T. Microbiol. Resour. Announc. 2020, 9, e01406–19.

65. Bolyen, E.; Rideout, J. R.; Dillon, M. R.; Bokulich, N. A.; Abnet, C. C.; Al-Ghalith, G. A.; Alexander, H.; Alm, E. J.; Arumugam, M.; Asnicar, F.; Bai, Y.; Bisanz, J. E.; Bittinger, K.; Brejnrod, A.; Brislawn, C. J.; Brown, C. T.; Callahan, B. J.; Caraballo-Rodríguez, A. M.; Chase, J.; Cope, E. K.; Da Silva, R.; Diener, C.; Dorrestein, P. C.; Douglas, G. M.; Durall, D. M.; Duvallet, C.; Edwardson, C. F.; Ernst, M.; Estaki, M.; Fouquier, J.; Gauglitz, J. M.; Gibbons, S. M.; Gibson, D. L.; Gonzalez, A.; Gorlick, K.; Guo, J.; Hillmann, B.; Holmes, S.; Holste, H.; Huttenhower, C.; Huttley, G. A.; Janssen, S.; Jarmusch, A. K.; Jiang, L.; Kaehler, B. D.; Kang, K. Bin; Keefe, C. R.; Keim, P.; Kelley, S. T.; Knights, D.; Koester, I.; Kosciolek, T.; Kreps, J.; Langille, M. G. I.; Lee, J.; Ley, R.; Liu, Y. X.; Loftfield, E.; Lozupone, C.; Maher, M.; Marotz, C.; Martin, B. D.; McDonald, D.; McIver, L. J.; Melnik, A. V.; Metcalf, J. L.; Morgan, S. C.; Morton, J. T.; Naimey, A. T.; Navas-Molina, J. A.; Nothias, L. F.; Orchanian, S. B.; Pearson, T.; Peoples, S. L.; Petras, D.; Preuss, M. L.; Pruesse, E.; Rasmussen, L. B.; Rivers, A.; Robeson, M. S.; Rosenthal, P.; Segata, N.; Shaffer, M.; Shiffer, A.; Sinha, R.; Song, S. J.; Spear, J. R.; Swafford, A. D.; Thompson, L. R.; Torres, P. J.; Trinh, P.; Tripathi, A.; Turnbaugh, P. J.; Ul-Hasan, S.; van der Hooft, J. J. J.; Vargas, F.; Vázquez-Baeza, Y.; Vogtmann, E.; von Hippel, M.; Walters, W.; Wan, Y.; Wang, M.; Warren, J.; Weber, K. C.; Williamson, C. H. D.; Willis, A. D.; Xu, Z. Z.; Zaneveld, J. R.; Zhang, Y.; Zhu, Q.; Knight, R.; Caporaso, J. G. Nat. Biotechnol. 2019, 37, 852–857.

66. McDonald, D.; Clemente, J. C.; Kuczynski, J.; Rideout, J. R.; Stombaugh, J.; Wendel, D.; Wilke, A.; Huse, S.; Hufnagle, J.; Meyer, F.; Knight, R.; Caporaso, J. G. Gigascience 2012, 464, 7.

67. Martin, M. EMBnet.journal 2011, 17, 10.

68. Callahan, B. J.; McMurdie, P. J.; Rosen, M. J.; Han, A. W.; Johnson, A. J. A.; Holmes, S. P. Nat. Methods 2016, 13, 581–583.

69. Quast, C.; Pruesse, E.; Yilmaz, P.; Gerken, J.; Schweer, T.; Yarza, P.; Peplies, J.; Glöckner, F. O. Nucleic Acids Res. 2013, 41.

70. Rognes, T.; Flouri, T.; Nichols, B.; Quince, C.; Mahé, F. PeerJ 2016, 2016, e2584.

71. Price, M. N.; Dehal, P. S.; Arkin, A. P. PLoS One 2010, 5, e9490.

72. Katoh, K.; Standley, D. M. Mol. Biol. Evol. 2013, 30, 772–780.

73. Lane, D. J. In Nucleic Acid Techniques in Bacterial Systematics; Stackebrandt, E.; Goodfellow, M.; Eds., John Wiley and Sons: New York, 1991; pp 115–175.

74. R Core Team. Vienna, Austria 2020.

75. Bisanz, J. E. 2018.

76. McMurdie, P. J.; Holmes, S. PLoS One 2013, 8, e61217.

77. Kandlikar, G. S.; Gold, Z. J.; Cowen, M. C.; Meyer, R. S.; Freise, A. C.; Kraft, N. J. B.; Moberg-Parker, J.; Sprague, J.; Kushner, D. J.; Curd, E. E. F1000Research 2018, 7, 1734.

78. Oksanen, J.; Blanchet, F. G.; Friendly, M.; Kindt, R.; Legendre, P.; McGlinn, D.; Minchin, P. R.; O’Hara, R. B.; Simpson, G. L.; Solymos, P.; Stevens, M. H. H.; Szoecs, E.; Wagner, H. 2019.

79. Wickham, H. Wiley Interdiscip. Rev. Comput. Stat. 2011, 3, 180–185.

