## Supporting Tables and Figures for "Secondary metabolites of *Bacillus subtilis* impact soil-derived semi-synthetic bacterial community assembly"

**Table S1.** Bacterial strains used in this study.

| Strains | Characteristics | Reference |
|---|---|---|
| P5_B1 | *B. subtilis* soil isolate from sampling site 55.788800, 12.558300 | [1] |
| DTUB55 | P5_B1 *sfp*::mls | [2] |
| DTUB79 | P5_B1 *srfAC*::Tn*10* (Spec$^R$) | [1] |
| DTUB102 | P5_B1 Δ*pksL* (Chl$^R$) | [2] |
| DTUB125 | P5_B1 Δ*ppsC* (Tet$^R$) | [2] |
| M5 | *Lysinibacillus fusiformis* | [3] |

**REFERENCES**


1. Thérien, M.; Kiesewalter, H. T.; Auria, E.; Charron-Lamoureux, V.; Wibowo, M.; Maróti, G.; Kovács, Á. T.; Beauregard, P. B. *Biofilm* **2020**, *2*, 100021.

2. Kiesewalter, H. T.; Lozano-Andrade, C. N.; Wibowo, M.; Strube, M. L.; Maróti, G.; Snyder, D.; Jørgensen, T. S.; Larsen, T. O.; Cooper, V. S.; Weber, T.; Kovács, Á. T. *bioRxiv* **2020**, 2020.08.05.238063.

3. Gallegos-Monterrosa, R.; Kankel, S.; Götze, S.; Barnett, R.; Stallforth, P.; Kovács, Á. T. *J. Bacteriol.* **2017**, *199*, e00204-17.

**Table S2.** 16S rRNA V3-V4 primer used in this study

| Name | Barcode | Primer |
|---|---|---|
| V3V4_forw_1 | TTTTAATC | CCTACGGGNGGCWGCAG |
| V3V4_rev_1 | TTTTAATC | GACTACHVGGGTATCTAATCC |
| V3V4_forw_2 | ATAATTAG | CCTACGGGNGGCWGCAG |
| V3V4_rev_2 | ATAATTAG | GACTACHVGGGTATCTAATCC |
| V3V4_forw_3 | ACCAAATT | CCTACGGGNGGCWGCAG |
| V3V4_rev_3 | ACCAAATT | GACTACHVGGGTATCTAATCC |
| V3V4_forw_4 | CTTATCAA | CCTACGGGNGGCWGCAG |
| V3V4_rev_4 | CTTATCAA | GACTACHVGGGTATCTAATCC |
| V3V4_forw_5 | TGATCATT | CCTACGGGNGGCWGCAG |
| V3V4_rev_5 | TGATCATT | GACTACHVGGGTATCTAATCC |
| V3V4_forw_6 | AGAATCTA | CCTACGGGNGGCWGCAG |
| V3V4_rev_6 | AGAATCTA | GACTACHVGGGTATCTAATCC |
| V3V4_forw_7 | TCAAGAAA | CCTACGGGNGGCWGCAG |
| V3V4_rev_7 | TCAAGAAA | GACTACHVGGGTATCTAATCC |
| V3V4_forw_8 | ATCGAAAT | CCTACGGGNGGCWGCAG |
| V3V4_rev_8 | ATCGAAAT | GACTACHVGGGTATCTAATCC |
| V3V4_forw_9 | ACATTTAC | CCTACGGGNGGCWGCAG |
| V3V4_rev_9 | ACATTTAC | GACTACHVGGGTATCTAATCC |
| V3V4_forw_10 | TAGAAAAC | CCTACGGGNGGCWGCAG |
| V3V4_rev_10 | TAGAAAAC | GACTACHVGGGTATCTAATCC |
| V3V4_forw_11 | TTATCACC | CCTACGGGNGGCWGCAG |
| V3V4_rev_11 | TTATCACC | GACTACHVGGGTATCTAATCC |
| V3V4_forw_12 | AATAGGGT | CCTACGGGNGGCWGCAG |
| V3V4_rev_12 | AATAGGGT | GACTACHVGGGTATCTAATCC |
| V3V4_forw_13 | ATTGCTGA | CCTACGGGNGGCWGCAG |
| V3V4_rev_13 | ATTGCTGA | GACTACHVGGGTATCTAATCC |
| V3V4_forw_14 | TGAGTTCT | CCTACGGGNGGCWGCAG |
| V3V4_rev_14 | TGAGTTCT | GACTACHVGGGTATCTAATCC |
| V3V4_forw_15 | GGCTATTT | CCTACGGGNGGCWGCAG |
| V3V4_rev_15 | GGCTATTT | GACTACHVGGGTATCTAATCC |
| V3V4_forw_16 | CAAGAGAT | CCTACGGGNGGCWGCAG |
| V3V4_rev_16 | CAAGAGAT | GACTACHVGGGTATCTAATCC |
| V3V4_forw_17 | GGAATACA | CCTACGGGNGGCWGCAG |
| V3V4_rev_17 | GGAATACA | GACTACHVGGGTATCTAATCC |
| V3V4_forw_18 | AAGGCAAT | CCTACGGGNGGCWGCAG |
| V3V4_rev_18 | AAGGCAAT | GACTACHVGGGTATCTAATCC |
| V3V4_forw_19 | ACAAAACG | CCTACGGGNGGCWGCAG |
| V3V4_rev_19 | ACAAAACG | GACTACHVGGGTATCTAATCC |
| V3V4_forw_21 | TTGAGTGA | CCTACGGGNGGCWGCAG |
| V3V4_rev_21 | TTGAGTGA | GACTACHVGGGTATCTAATCC |
| V3V4_forw_22 | GCTTCTGA | CCTACGGGNGGCWGCAG |
| V3V4_rev_22 | GCTTCTGA | GACTACHVGGGTATCTAATCC |
| V3V4_forw_23 | GGCAAGAT | CCTACGGGNGGCWGCAG |
| V3V4_rev_23 | GGCAAGAT | GACTACHVGGGTATCTAATCC |

| | | |
|---|---|---|
| V3V4_forw_24 | GTGCTTTC | CCTACGGGNGGCWGCAG |
| V3V4_rev_24 | GTGCTTTC | GACTACHVGGGTATCTAATCC |
| V3V4_forw_25 | ACACACTG | CCTACGGGNGGCWGCAG |
| V3V4_rev_25 | ACACACTG | GACTACHVGGGTATCTAATCC |
| V3V4_forw_26 | CGATTCTG | CCTACGGGNGGCWGCAG |
| V3V4_rev_26 | CGATTCTG | GACTACHVGGGTATCTAATCC |
| V3V4_forw_27 | GCAGAGTT | CCTACGGGNGGCWGCAG |
| V3V4_rev_27 | GCAGAGTT | GACTACHVGGGTATCTAATCC |
| V3V4_forw_28 | CGTCCTAT | CCTACGGGNGGCWGCAG |
| V3V4_rev_28 | CGTCCTAT | GACTACHVGGGTATCTAATCC |
| V3V4_forw_30 | GCTTGGTT | CCTACGGGNGGCWGCAG |
| V3V4_rev_30 | GCTTGGTT | GACTACHVGGGTATCTAATCC |
| V3V4_forw_31 | ACAGGCTT | CCTACGGGNGGCWGCAG |
| V3V4_rev_31 | ACAGGCTT | GACTACHVGGGTATCTAATCC |
| V3V4_forw_33 | TGACGCTT | CCTACGGGNGGCWGCAG |
| V3V4_rev_33 | TGACGCTT | GACTACHVGGGTATCTAATCC |

**Table S3.** Number of reads per sample.

| **Sample** | **Number of reads** |
|---|---|
| Soil 1 | 751 |
| Soil 2 | 12,660 |
| Pre 1 | 20,440 |
| Pre 2 | 17,787 |
| Pre 3 | 15,471 |
| Pre 4 | 34,802 |
| Control 1 | 4,649 |
| Control 2 | 20,846 |
| Control 3 | 16,229 |
| Control 4 | 16,556 |
| WT 1 | 12,436 |
| WT 2 | 18,554 |
| WT 3 | 18,182 |
| WT 4 | 11,964 |
| *sfp* 1 | 13,354 |
| *sfp* 2 | 16,737 |
| *sfp* 3 | 12,472 |
| *sfp* 4 | 21,421 |
| *srfAC* 1 | 7,047 |
| *srfAC* 2 | 1,184 |
| *srfAC* 3 | 3,563 |
| *srfAC* 4 | 7,663 |
| Δ*pksL* 1 | 11,017 |
| Δ*pksL* 2 | 2,115 |
| Δ*pksL* 3 | 7,063 |
| Δ*pksL* 4 | 5,187 |
| Δ*ppsC* 1 | 9,443 |
| Δ*ppsC* 2 | 10,823 |
| Δ*ppsC* 3 | 9,947 |
| Δ*ppsC* 4 | 2,112 |

### Supporting figures

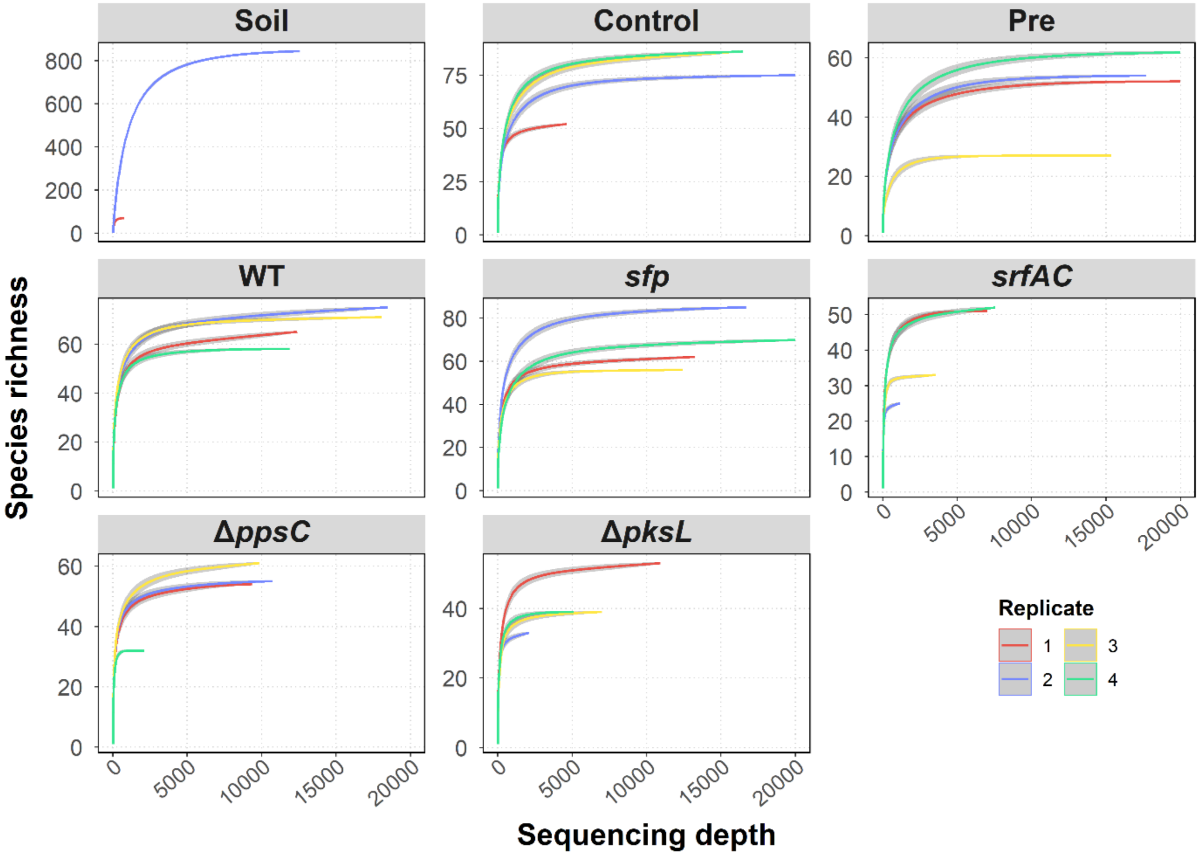


**Figure S1.** Rarefaction curves for each assay and replicate. Species richness is visualised for sampling depths from 0 to 20,000.

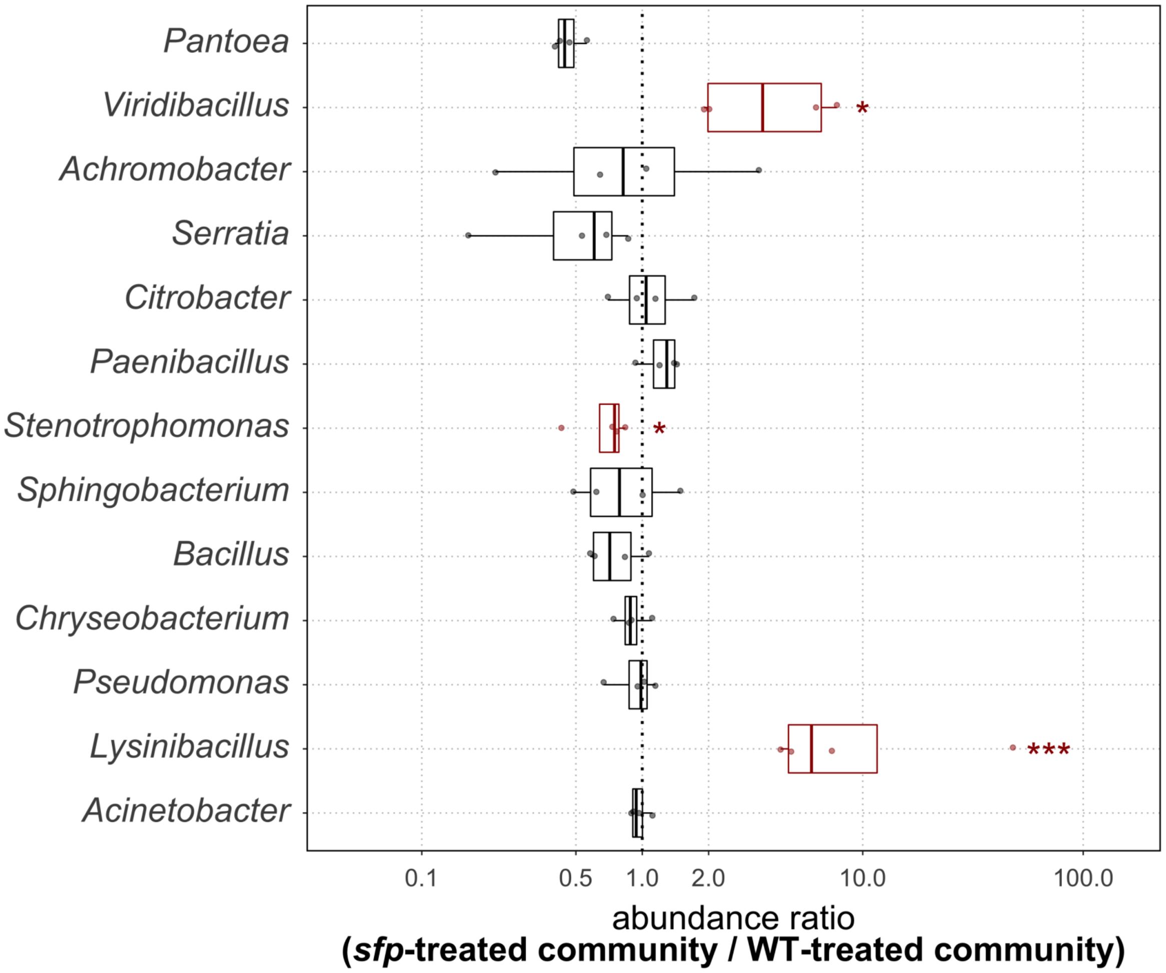


**Figure S2.** Abundance ratios for each genus and replicate (points) in the *sfp*-treated community compared to the WT-treated community. Statistical significance is defined as P ≤ 0.05 (*) and P ≤ 0.001 (***).

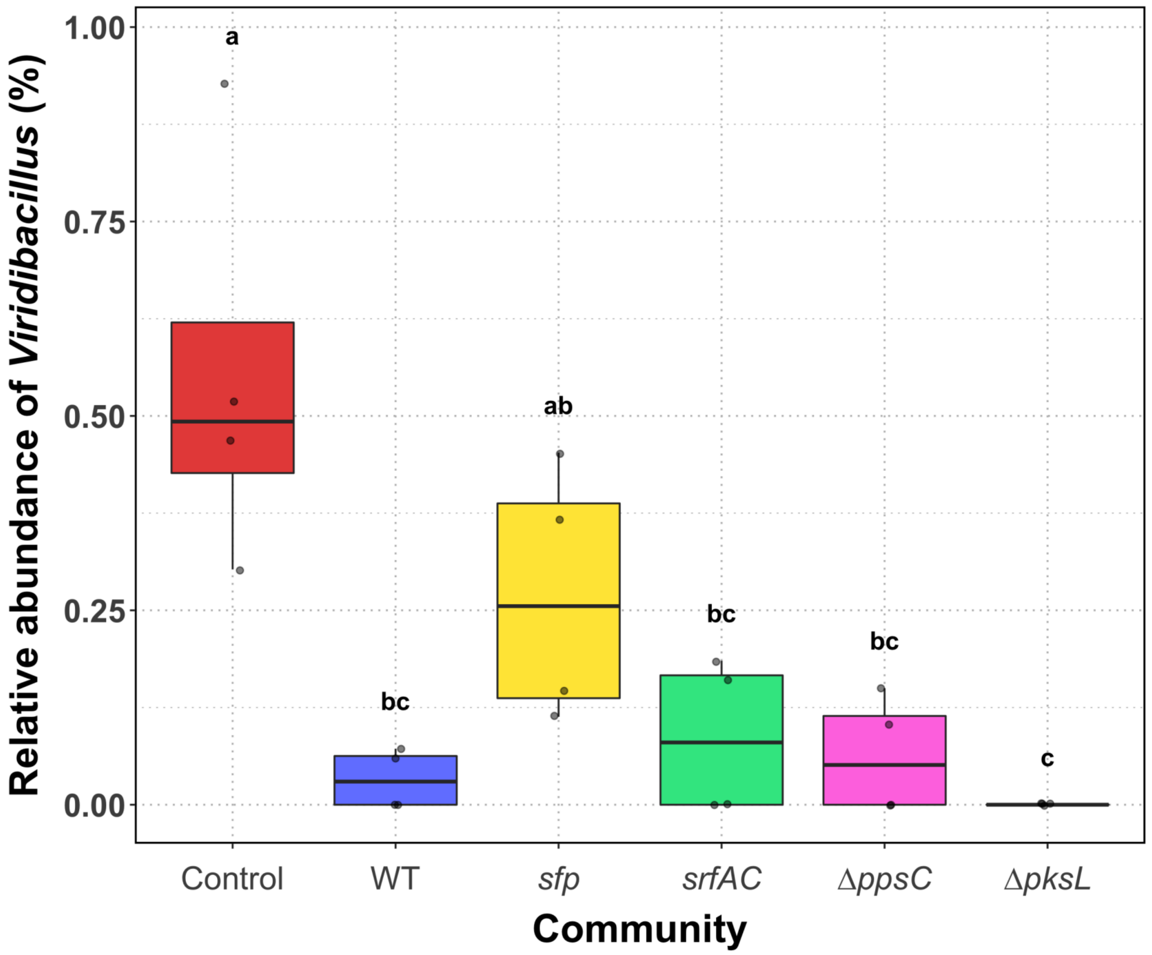


**Figure S3.** Abundance of *Viridibacillus* in the untreated mock communities (Control) and treated mock communities with either *B. subtilis* wild type (WT), NRPs deficient strain (*sfp*), surfactin mutant (*srfAC*), plipastatin mutant (Δ*ppsC*) or bacillaene mutant (Δ*pksL*) and co-cultivated for 48 h. Points represent the abundance in each replicate. Treatments with varying letters are significantly different ($P < 0.05$).

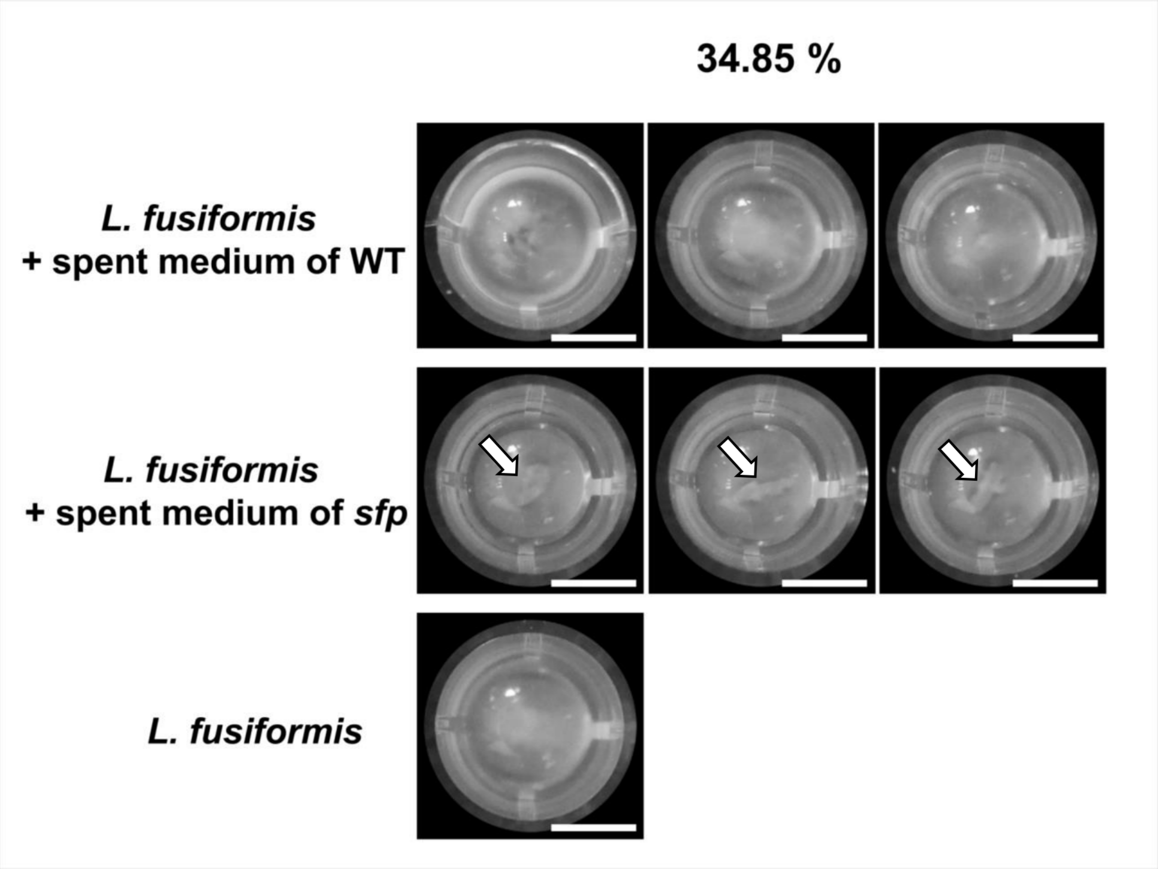


**Figure S4.** Different levels of aggregation of *L. fusiformis* M5 after exposure to spent medium of *B. subtilis* WT and *sfp* or without treatment. Solid aggregates in assays supplemented with spent medium of *sfp* are marked with arrows. Scale bars indicate 3 mm.

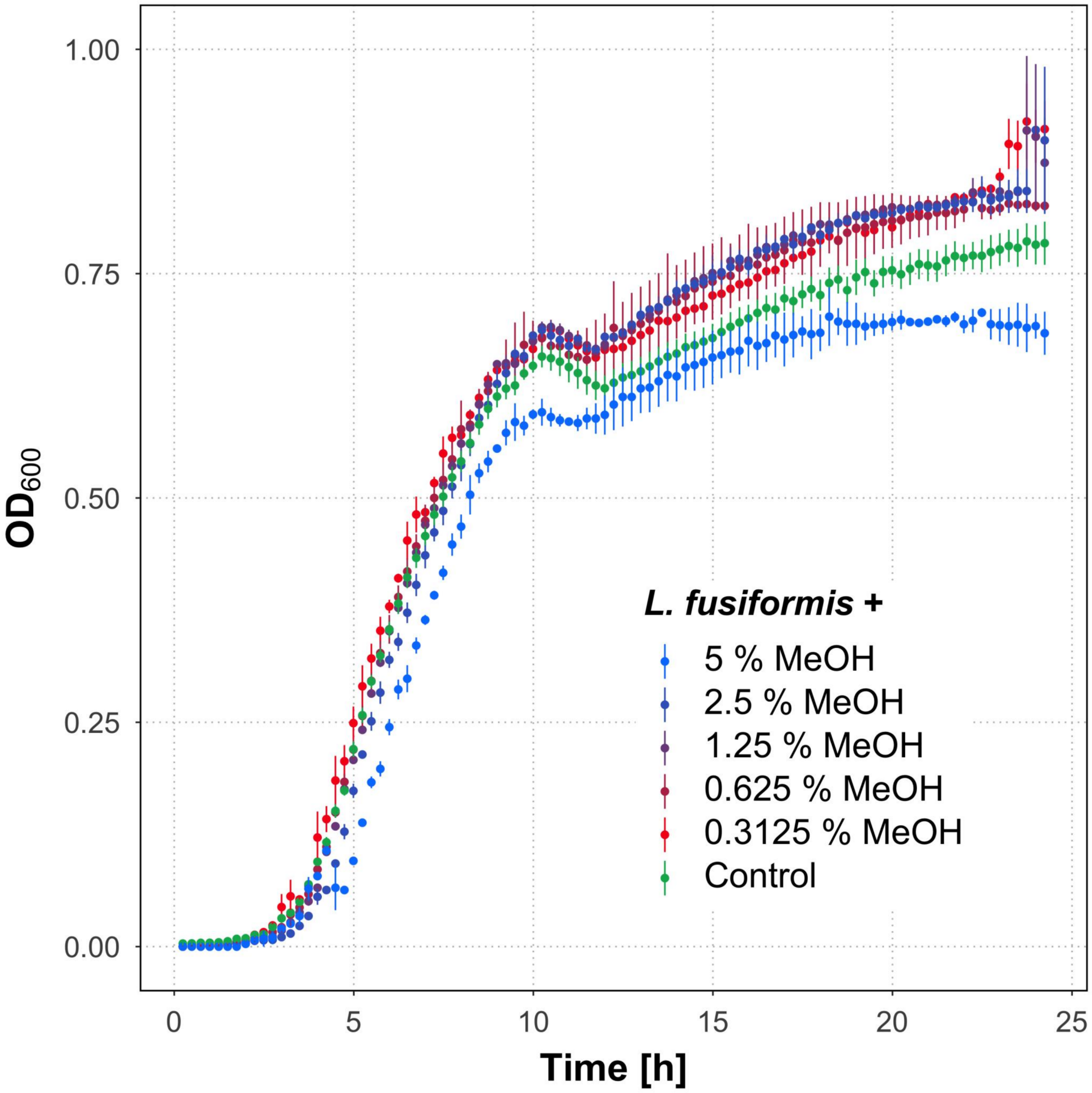


**Figure S5.** Growth curves of *L. fusiformis* M5 exposed to different concentrations of the solvent MeOH and without treatment (control). Error bars represent the standard error. N = 6 (control assay), N = 2 (MeOH-treated assays)
